# Role of CaMKIIa reticular neurons of caudal medulla in control of posture

**DOI:** 10.1101/2025.03.17.643745

**Authors:** Pavel V. Zelenin, Vladmir F. Lyalka, Shih-Hsin Chang, Francois Lallemend, Tatiana G. Deliagina, Li-Ju Hsu

**Affiliations:** Department of Neuroscience, Karolinska Institutet, 171 77 Stockholm, Sweden

## Abstract

Terrestrial quadrupeds actively stabilize dorsal-side-up orientation of the body in space due to activity of the postural control system. Supraspinal influences, including those from the reticular formation, play a crucial role in the operation of this system. However, the role of specific molecularly identified populations of reticular neurons in control of posture remains unknown. The aim of the present study was to reveal the role of CaMKIIa reticular neurons (CaMKIIa-RNs) located in the caudal medulla in control of posture. For this purpose, the effects of unilateral chemogenetic activation/inactivation of CaMKIIα-RNs on different aspects of postural control were studied in mice. It was found that unilateral activation of CaMKIIa-RNs evoked ipsilateral roll tilt of the head and trunk, caused by flexion/adduction of the ipsilateral limbs and extension/abduction of the contralateral limbs. The body roll tilt was actively stabilized on the tilting platform and maintained during walking. Unilateral inactivation of CaMKIIa-RNs evoked the opposite effects. Histological analyses showed that the population of CaMKIIa-RNs in the caudal medulla contains reticulospinal neurons that project to the spinal cord mainly through ipsilateral lateral funiculus and terminate in the intermediate area of the gray matter. We demonstrated that although the population of CaMKIIa-RNs contains both excitatory and inhibitory neurons, the excitatory ones dominate. Thus, CaMKIIa-RNs located in the caudal medulla play a crucial role for maintenance of the dorsal-side-up body orientation in different environments. Left/right symmetry and asymmetry in activity of CaMKIIa-RNs allows animals to maintain dorsal-side-up body orientation on horizontal and laterally inclined surfaces, respectively.

## INTRODUCTION

The maintenance of the basic body posture (upright in humans and a dorsal-side-up in terrestrial quadrupeds) is a vital motor function. Any deviation from this orientation induced by external forces triggers an automatic postural response – a corrective movement – aimed at restoring the initial orientation. Also, both humans and animals can specifically change the body configuration in context of different motor behaviors.

The basic postural networks reside in the brainstem, cerebellum, and spinal cord (Musienko et al., 2008). Supraspinal networks play a crucial role in control of posture. Distortions in activity of descending systems forming the output of the postural networks – potentially caused by spinal cord injury or by neurological diseases – results in severe postural dysfunctions (Lyalka et al., 2005, 2009, 2011; Macpherson et al., 1997; Macpherson and Fung, 1999; Rossignol et al., 1999, 2002; Barbeau et al., 2002; Chvatal et al., 2013).

It was suggested that there are two types of posture-related supraspinal influences: first, phasic postural commands (Karayannidou et al., 2008; Zelenin et al., 2010; Stapley and Drew 2009; Matsuyama and Drew, 2000) contributing to generation of postural corrections and, second, tonic drive activating spinal postural networks (Musienko et al., 2010; Lyalka et al., 2011; Zelenin et al., 2013).

Neurons of reticular formation of the brainstem are important elements of supraspinal postural networks. They form a number of reticular nuclei from which originates the phylogenetically oldest descending system—the reticulospinal one. In the lamprey (a lower vertebrate animal), reticulospinal system is the only developed supraspinal system and it plays a key role in control of posture. It was demonstrated that any deviation from the stabilized dorsal side up body orientation in the lamprey leads to activation of a specific population of reticulospinal neurons (Deliagina et al., 2014). Each of the neurons in this population activates a specific motor synergy and collectively, the activated reticulospinal neurons evoke the motor output necessary for the postural correction (Zelenin et.al., 2007). Also, it was demonstrated that left/right asymmetry in activity of RS neurons shifts the set-point of postural control system leading to stabilization of a new orientation of the body in space (Deliagina et al., 2014).

A number of evidence indicates that reticulospinal neurons in higher vertebrates play similar functional roles in control of posture. It was demonstrated in cats that reticulospinal neurons in the pontomedullary reticular formation transmit phasic postural commands for generation of postural corrections caused by an unexpected drop of support under one of the limbs (Stapley and Drew, 2009). Previously, we demonstrated that in rabbits, binaural galvanic vestibular stimulation (GVS) evokes lateral body sway that is actively stabilized indicating that GVS shifts the set-point of the postural control system (Beloozerova et al., 2003; Hsu et al., 2012). Since GVS evokes strong asymmetry in activity of vestibular afferents (Goldberg et al., 1984; Minor and Goldberg, 1991) and reticulospinal neurons receive substantial vestibular input (Wilson and Melvill Jones, 1979), one can suggest that left/right asymmetry in activity of reticulospinal neurons contributes to the shift of the set-point of the postural system. It was also demonstrated that microstimulation of specific sites within the pontomedullary reticular formation activates different motor synergies (involving muscles of both left and right limbs), which can contribute to generation of postural corrections as well as to changes of the body configuration in context of specific motor behaviors while stimulation of other sites evokes bilateral augmentation or suppression of the muscle tone (Takikusaki et al., 2016).

Although it is documented that neurons of the pontomedullary reticular formation contribute to control of posture, the role of different molecularly identified reticular neurons located in different parts of reticular formation in control of particular aspects of posture, such as the body orientation in space, efficacy of postural corrections, specific changes of the body configuration, remains unknown.

Recent advances in genetics have inspired numerous studies striving to identify the functional role of neurons, and in particular reticular ones, in control of movements based on transcription factor expression. It was demonstrated that Vglut2 reticular neurons located in the lateral paragigantocellular nucleus are implicated in initiation of locomotion (Hsu et al., 2023; Capelli et al., 2017), while Chx10 V2a reticular neurons located in the gigantocellular reticular nucleus cause locomotor stop and left/right asymmetry in their activity evokes lateral turn (Cregg et al., 2020; Usseglio et al., 2020; Bouvier et al., 2015).

In the present study, we revealed a specific population of reticular neurons, the Calcium– calmodulin-dependent protein kinase II alpha expressing reticular neurons (CaMKIIa-RNs), located in the caudal medulla that control the body orientation in the transverse plane in mice. We demonstrated that excitatory neurons dominate in the CaMKIIa-RNs population and that it contains reticulospinal neurons.

## RESULTS

To investigate the role of CaMKIIa-RNs in control of posture, we used the chemogenetic approach. *First,* excitatory (hM3Dq) or inhibitory (hM4Di) DREADDs were expressed unilaterally in CaMKIIa-RNs of the caudal medulla in mice (Fig. 1**A**). *Second*, the mice performed each of four basic motor behaviors (standing on a horizontal surface, postural corrections on a tilting platform, forward locomotion, and righting) before and during unilateral activation/inactivation of CaMKIIa-RNs caused by CNO injection. Motor performance before and during activation/inactivation of CaMKIIa-RNs were analyzed and compared. In the following text, terms “ipsilateral” and “contralateral” are used to indicate, respectively, the ipsilateral and contralateral side in relation to the side of the virus injection.

**Figure 1.**
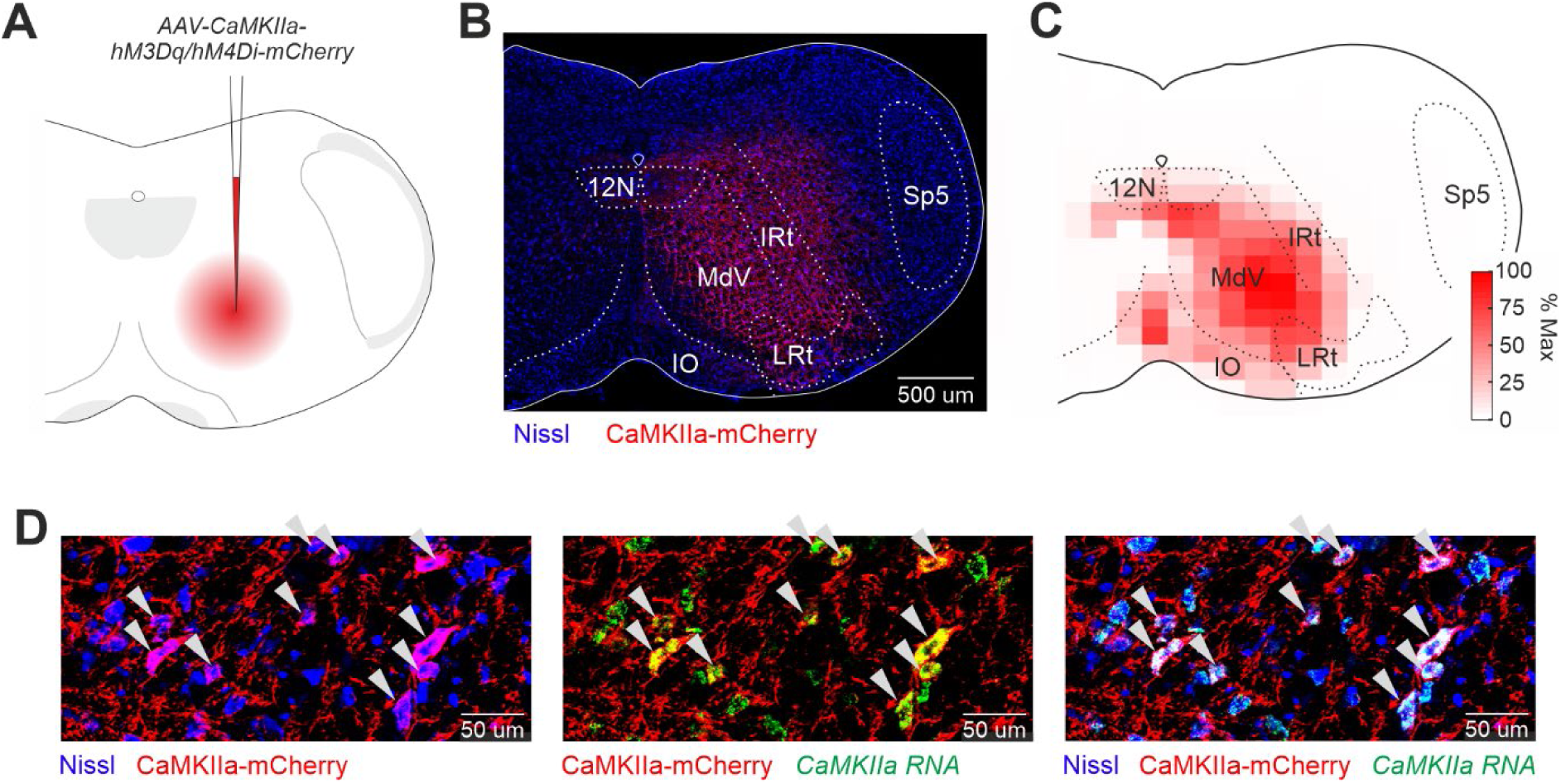
Unilateral expression of DREADDS in CaMKIIa-RNs of the caudal medulla. **A.** A schematic drawing of the AAV-CaMKIIa-hM3Dq/hM4Di-mCherry injection site. **B**. A representative example of unilateral infection of CaMKIIa-RNs with AAV-CaMKIIa-hM3Dq-mCherry. **C**. A heatmap showing the average extent of the infected area (*N*=13). **D**. Combination of Nissl staining with immunochemistry for mCherry and *in situ* hybridization for CaMKIIa mRNA showing that the infected cells are CaMKIIa^+^ neurons (indicated by white arrowheads). *MdV*, the medullary reticular nucleus ventral part; *IRt*, the intermediate reticular nucleus; *LRt*, the lateral reticular nucleus; *IO*, the inferior olive: *12N*, the hypoglossal nucleus; *Sp5*, the spinal trigeminal nucleus.

A representative example of an infected area (CaMKIIa-RNs expressing DREADDs) is shown in Fig. 1**B**. The infected area covered mainly the medullary reticular nucleus ventral part (MdV) and the intermediate reticular nucleus (IRt), as seen in the heatmap of the intensity of the mCherry fluorescence averaged across all animals (Fig. 1**C**). The specificity of DREADDs expression in CaMKIIa-RNs was confirmed by co-localization of CaMKIIa mRNA and mCherry (cells indicated by white arrowheads in Fig. 1**D**).

### Left-right asymmetry in activity of CaMKIIa-RNs evokes the body roll tilt in the animal standing on a horizontal surface

To reveal changes in the basic body posture of the mouse standing on a horizontal surface, its frontal and rear views (Fig. 2**A**,**D**) as well as the view from below (Fig. 2**G**) were video recorded before and during unilateral activation/inactivation of CaMKIIa-RNs (respectively, *Control* and *CNO* in Fig. 2**A**,**D**,**G**).

**Figure 2.**
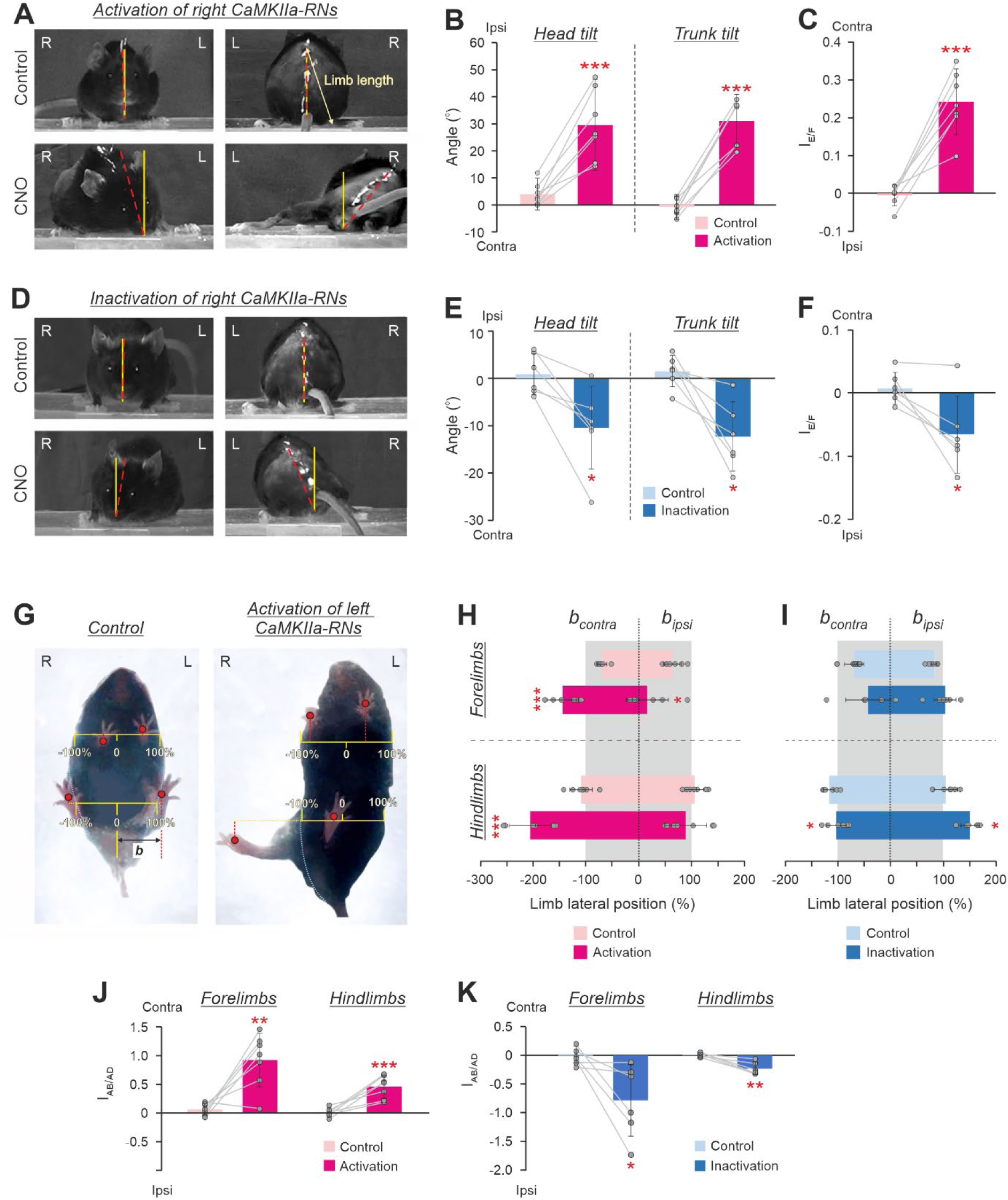
Effects of unilateral activation/inactivation of CaMKIIa-RNs on the body posture in mice standing on a horizontal surface. **A,D**. The front and rear views of mice before (*Control*) and during activation (**A**, *CNO*) and inactivation (**D**, *CNO*) of the right CaMKIIa-RNs. The solid yellow line indicates the vertical. The dashed red line indicates the dorso-ventral axis of the head or trunk. **B,E**. Values of the head and the trunk roll tilt angles in individual animals, as well as corresponding mean ± SD values, before (*Control*) and during unilateral activation (**B)** and inactivation (**E)** of CaMKIIa-RNs. **C**,**F**. Values of the extension/flexion asymmetry index (*I_E/F_*) in individual animals, as well as the corresponding mean ± SD values, before (*Control*) and during unilateral activation (**C**) and inactivation (**F**) of CaMKIIa-RNs. **G.** The view from below of a mouse before (*Control*) and during activation of the left CaMKIIa-RNs. **H,I.** The lateral positions of the contralateral and the ipsilateral limbs in individual animals as well as the corresponding mean ± SD values, before (*Control*) and during unilateral activation (**H**) and inactivation (**I**) of CaMKIIa-RNs. The lateral position is measured in percents of the corresponding half body width. **J,K**. The abduction/adduction asymmetry index of the forelimbs and hindlimbs (*I_AB/AD_*) in individual animals, as well as the corresponding mean ± SD values, before (*Control*) and during unilateral activation and inactivation of CaMKIIa-RNs. In **B,C,H**: *N* = 7. In **E,F,I**: *N* = 6. In **J,K**: *N* = 7 and *N* = 6 for activation and inactivation of CaMKIIa-RNs, respectively. L and R, left and right, respectively. *FL* and *HL*, forelimb and hindlimb, respectively. *Ipsi* and *Contra,* ipsilateral and contralateral in relation to the virus injection side, respectively. Indication of significance level: * 0.01< *p* < 0.05, ** 0.001 < *p* < 0.01, *** *p* < 0.001.

We found that, while before CNO administration animals maintained the dorsal side-up orientation of the head and trunk (*Control* in Fig. 2**A**,**D**), injection of CNO caused a gradually developing roll tilt of the head and trunk that reached its maximal expression in about 40 minutes after CNO injection (*CNO* in Fig. 2**A**,**D**). The maximal effect lasted for about 1-1.5 hour, then gradually decreased, and disappeared completely in 3-4 hours after the injection.

We found that unilateral activation of CaMKIIa-RNs evoked ipsilateral (in relation to the side of the activated neurons) roll tilt of the head and trunk (compare *Control* and *CNO* in Fig. 2**A**). On average, a significant increase in the values of the ipsilateral head and trunk tilt angles during unilateral activation of CaMKIIa-RNs, as compared to those in control were observed (30±17° *vs* 4±6° for the head tilt angle and 31±10° *vs* -1±5° for the trunk tilt angle; paired *t* test, *p* = 9×10^-4^ and *p* = 6×10^-5^, respectively; Fig. 2**B**).

The ipsilateral roll tilt of the trunk was caused by a change in configuration of the left and right limbs as well as in their position in relation to the trunk. To reveal asymmetry in the configuration of the ipsilateral and contralateral limbs caused by unilateral activation of CaMKIIa-RNs, we compared the limbs length as well as their lateral positions in relation to the trunk before and after CNO injection.

To estimate the asymmetry in the limbs length, we calculated the extension/flexion asymmetry index for hindlimbs during standing based on the difference between lengths of the contralateral and ipsilateral limb (see Methods for details). We found that in control, the asymmetry index was close to 0 (-0.01±0.03), indicating that the lengths of the left and right limbs were almost equal (Fig. 2**C**). By contrast, during unilateral activation of CaMKIIa-RNs, the asymmetry index was positive (0.24±0.08), indicating that the contralateral limb was longer (more extended) than the ipsilateral one (Fig. 2**C**). The difference between the mean value of the extension/flexion asymmetry index of hindlimbs during unilateral activation of CaMKIIa-RNs and that observed in control was statistically significant (paired *t* test, *p* = 3×10^-4^; Fig. 2**C**). Thus, unilateral activation of CaMKIIa-RNs evoked asymmetry in configuration of the ipsilateral and contralateral hindlimbs – the contralateral limb became more extended than the ipsilateral one.

To estimate the asymmetry in the lateral positions of the limbs in relation to the trunk, we calculated the abduction/adduction asymmetry index for forelimbs and hindlimbs (see Methods for details). In control, the lateral positions of the left and right limbs in relation to the trunk midline, were almost symmetrical (Fig. 2**G**, left panel) and the values of the abduction/adduction asymmetry index for both forelimbs and hindlimbs were close to 0 (Fig. 2**J**). By contrast, during unilateral activation of CaMKIIa-RNs, the lateral positions of the left and right limbs were highly asymmetrical: both the ipsilateral forelimb and hindlimb were closer to the trunk midline than the contralateral limbs (Fig. 2**G**, right panel). On average, the lateral positions of the contralateral forelimb and hindlimb (expressed in percent of the half of the corresponding body width) were significantly larger than in control (respectively, - 144±30% *vs* -70±9% and -206±39% *vs* -110±21%; paired *t* test, *p* = 6×10^-4^ and *p* = 0.001, Fig. 2**H**). By contrast, the lateral positions of the ipsilateral forelimb as well as hindlimb were on average smaller than in control and significant only for forelimb (respectively, 16±40% and 64±17%*vs* 89±40% and 106±18%; paired *t* test, *p* = 0.03 and *p* = 0.34; Fig. 2**H**). The asymmetry in the lateral positions of the left and right limbs was also reflected in the values of the abduction/adduction asymmetry index that were significantly higher than those in control for both fore- and hindlimbs (respectively, 0.92±0.47 *vs* 0.07±0.11 and 0.46±0.20 *vs* 0.01±0.08, paired *t* test, *p* = 0.002 and *p* = 7×10^-4^; Fig. 2**J**). Note that after CNO the values of the abduction/adduction asymmetry indexes were positive indicating that contralateral limbs were more abducted than ipsilateral ones. Thus, these results suggest that unilateral activation of CaMKIIa-RNs evokes abduction of the contralateral and adduction of the ipsilateral limbs.

Unilateral inactivation of CaMKIIa-RNs evoked effects opposite to those observed during unilateral CaMKIIa-RNs activation. It resulted in the contralateral (in relation to the inactivated neurons) roll tilt of the head and trunk (Fig. 2**D,E**) caused by extension and abduction of ipsilateral limbs as well as by flexion and adduction of the contralateral limbs (Fig. 2**F,I**). On average, parameters such as head and trunk roll angles and the limbs extension/flexion asymmetry index were significantly different compared to those in control (head angle: -10±9° after CNO *vs* 1±4° in control; trunk angle: -12±7° after CNO *vs* 1±3°; extension/flexion asymmetry index: -0.06±0.06 after CNO *vs* 0.01±0.24 in control; paired *t* test, *p* = 9×10⁻⁴, *p* = 6×10⁻⁵, *p* = 0.02, respectively; Fig. **2E,F**). However, the absolute values of their changes were almost four times smaller than during unilateral activation of CaMKIIa-RNs (compare the corresponding parameters in **E** and **B**, **F** and **C** in Fig. 2). Also, abduction of the ipsilateral limbs and adduction of the contralateral limbs were much weaker expressed during unilateral CaMKIIa-RNs inactivation (Fig. 2**I**) as compared to those observed during unilateral activation (Fig. 2**H**). Nevertheless values of these measures were significant as compared to control for the hindlimbs (respectively, 153±17% *vs* 110 ± 18% and -99±21% *vs* -114±14%; paired *t* test, *p* = 0.03 and *p* = 0.03, Fig. 2**I**). Despite the weaker effect, inactivation of CaMKIIa-RNs evoked a significant asymmetry in the lateral position of the left and right limbs. During inactivation, the values of the abduction/addiction asymmetry index for both fore- and hindlimbs differed significantly from those in control (respectively, -0.79±0.63 *vs* 0.03±0.15 and -0.24±0.11 *vs* 0.01±0.04, paired *t* test, *p* = 0.003 and *p* = 0.003; Fig. 2**K**) and had negative values indicating that the ipsilateral limbs were more abducted than contralateral ones.

In summary, a left-right asymmetry in activity of CaMKIIa-RNs located in MdV-IRt area evoked the body roll tilt toward more active (dominant) sub-population of CaMKIIa-RNs. The body roll tilt was caused by flexion and adduction of limbs on the dominant side and by simultaneous extension and abduction of the limbs on the opposite side. The effects of activation of the CaMKIIa-RNs were stronger than those of inactivation.

### The roll tilt caused by asymmetry in activity of CaMKIIa-RNs is actively stabilized during standing on the tilting platform

To find out whether the body roll tilt evoked by unilateral activation/inactivation of CaMKIIa-RNs during standing on a horizontal surface is actively stabilized, we analyzed postural corrections of mice standing on the platform subjected to lateral tilts (Fig. 3).

**Figure 3.**
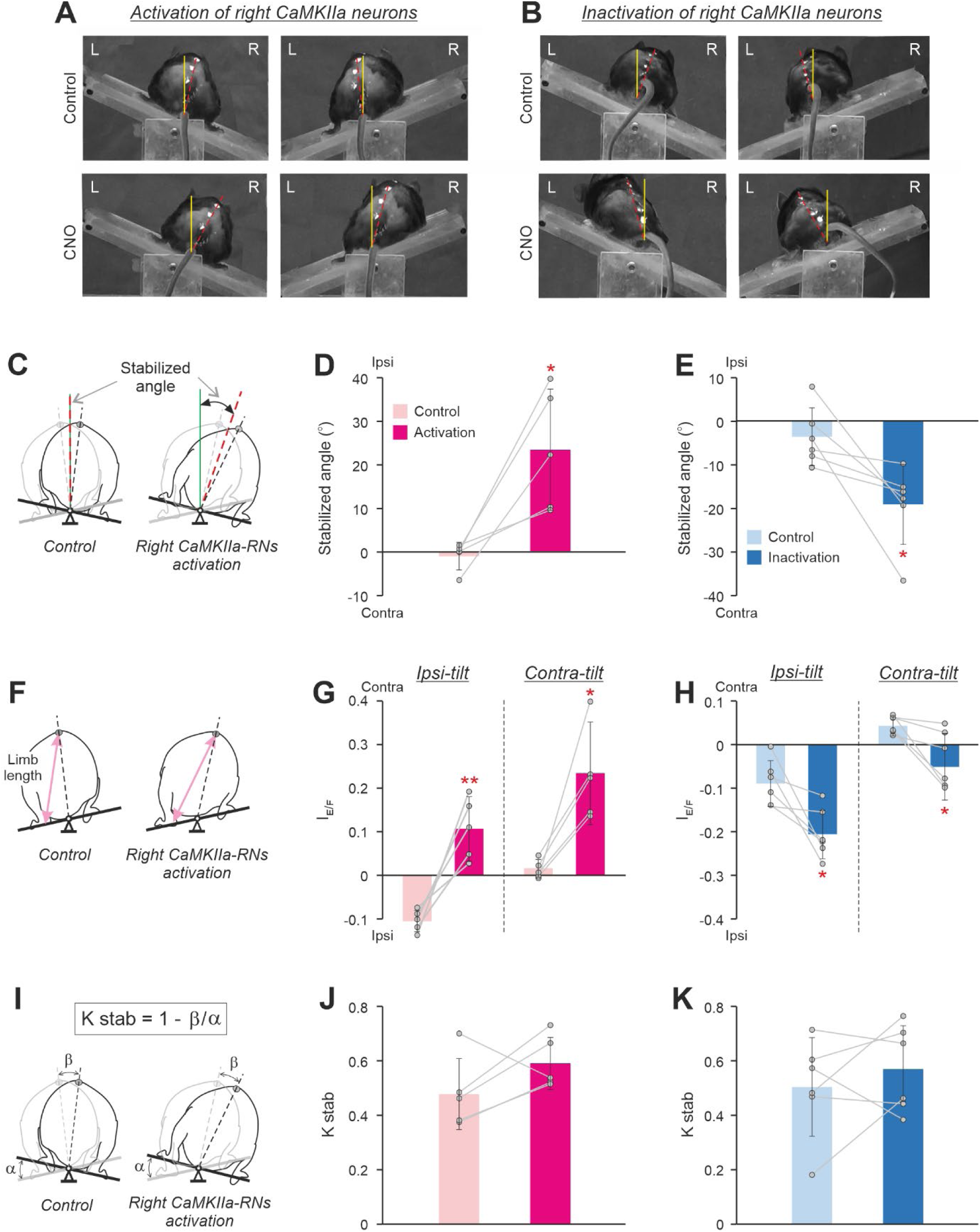
Effects of unilateral activation/inactivation of CaMKIIa-RNs on postural corrections caused by lateral tilts of the platform. **A,B.** The rear views of mice standing on the platform tilted to the right (left panels) and to the left (right panels) before (*Control*) and after activation (**A**, *CNO*) and inactivation (**B**, *CNO*) of the right CaMKIIa-RNs. Designations as in Fig. 2A**,B**. **C.** Schematic drawings indicating the “*Stabilized angle*” on the tilting platform before (*Control*) and during activation of the right CaMKIIa-RNs (see Methods for details). The black and gray dashed lines indicate the dorso-ventral axis of the trunk when the animal is standing on the platform tilted to the right and to the left, respectively. The green solid line indicates the vertical. The dashed crimson line indicates the bisector of the angle formed by the dorso-ventral axis of the trunk at two conditions: when the animal is standing on the platform tilted to the left and when it standing on the platform tilted to the right. **D,E.** Values of the stabilized angle in individual animals, as well as the corresponding mean ± SD values, before (*Control*) and during unilateral activation (**D)** and inactivation (**E)** of CaMKIIa-RNs. **F.** Schematic drawings indicating left hindlimb lengths (shown by the pink line with arrows) during standing on the platform tilted to the left before (*Control*) and during activation of the right CaMKIIa-RNs. **G,H.** Values of the extension/flexion asymmetry index during standing (*I_E/F_*) in individual animals, as well as the corresponding mean ± SD values, during standing, respectively, on the ipsilaterally and contralaterally tilted platform (*Ipsi-tilt* and *Contra-tilt*, respectively) before (*Control*) and after unilateral activation (**G**) and inactivation (**H)** of the CaMKIIa-RNs. **I.** Schematic drawings explaining the estimation of the efficacy of postural corrections (*KSTAB*). α, the amplitude of the platform tilt; β, the amplitude of the dorso-ventral axis of the trunk tilt during standing on the tilting platform. **J,K.** Values of the Kstab in individual animals, as well as the corresponding mean ± SD values, before (*Control*) and during unilateral activation (**J)** and inactivation (**K)** of CaMKIIa-RNs. In **D,G,J**: *N* = 5. In **E,H,K**: *N* = 6. Abbreviations as in Fig. 2.

Before CNO injection, a lateral tilt of the supporting platform evoked extension of limbs on the side of the tilt and simultaneous flexion of the contralateral limbs (*Control* in Fig. 3**A**,**B**) leading to a displacement of the dorso-ventral axis of the trunk (indicated by the red dashed line in Fig. 3**A**,**B**) towards the vertical (indicated by the solid yellow line in Fig. 3**A**,**B**). However, as in all tested terrestrial quadrupeds (Deliagina et al., 2000, 2006; Beloozerova et al., 2003), the postural corrections in mice did not fully compensate the distortion of the trunk orientation caused by the platform tilt, and after their execution, the dorso-ventral axis of the trunk was still deviated from the vertical (*Control* in Fig. 3**A**,**B**; Zelenin et al., 2021).

The orientation of the trunk stabilized on the tilting platform was characterized by a “Stabilized angle” (see Methods for details; Fig. 3C). Before CNO injection, the stabilized angle was close to zero suggesting that the animal stabilized close to the dorsal-side-up trunk orientation (*Control* in Fig. 3**D**,**E**). We found that during unilateral activation of CaMKIIa-RNs, the value of the stabilized angle was positive and significantly larger than in control (24±14° *vs* –1±3°, paired *t* test, *p* = 0.02; Fig 3**D**) suggesting that the animal stabilized the body orientation with an ipsilateral roll tilt. By contrast, during unilateral inactivation, the stabilized angle value was negative and significantly different from that in control (–19±9° *vs* –4±7°, paired *t* test, *p* = 0.03; Fig. 3**E**) suggesting that the animal stabilized the body orientation with a contralateral roll tilt.

The change of the stabilized trunk orientation was caused by changes in configurations of the left and right limbs performing the corrective movements. In control, at condition when the contralateral and ipsilateral limb were standing on the side of the tilt, the averaged extension/flexion asymmetry index had, respectively, positive and negative values, indicating that the length of the limb on the side of the tilt was longer than the length of the opposite limb (*Control* in Fig. 3**G,H**). The increase in the limb length (the limb extension) on the side of the tilt and the simultaneous decrease in the length (flexion) of the opposite limb moved the dorso-ventral axis of the trunk toward the vertical. We found that during unilateral activation of CaMKIIa-RNs, the mean value of the extension/flexion asymmetry index was positive during both ipsilateral and contralateral tilts (Fig. 3**G**). Also, the values of the extension/flexion asymmetry index differed significantly from those in control (0.11±0.07 vs -0.10±0.02 for the ipsilateral tilt and 0.24±0.12 *vs* 0.01±0.02 for the contralateral tilt; paired *t* test, *p* = 0.005 and *p* = 0.02, respectively; Fig. 3**G**).

These results suggest that asymmetry in the hindlimbs length evoked by unilateral activation of CaMKIIa-RNs in animals standing on a horizontal surface was maintained during postural corrections. During both ipsilateral and contralateral tilt, the length of the contralateral limb was larger than that of the ipsilateral one. Thus, contralateral and ipsilateral limb performed corrective movements with more extended and flexed configuration, respectively, as compared to control. This led to the positive value of the stabilized angle, reflecting stabilization of the trunk orientation with the ipsilateral roll tilt.

We found that unilateral inactivation of CaMKIIa-RNs led to the opposite effects (Fig. 3**H**). During both ipsilateral and contralateral tilts, the mean values of the extension/flexion asymmetry index were negative and significantly different from corresponding values in control (ipsilateral tilt: -0.20±0.06 during CNO *vs* -0.09±0.05 in control, paired *t* test, *p* = 0.03; contralateral tilt: -0.05±0.08 during CNO *vs* 0.04±0.02 in control, paired *t* test, *p* = 0.03). Thus, the ipsilateral limb performed corrective movements with more extended configuration, while the contralateral limb performed postural corrections with more flexed configuration as compared to those in control. This led to the negative value of the stabilized angle, reflecting stabilization of the trunk orientation with the contralateral roll tilt (Fig. 3**E**).

To reveal possible effects of unilateral activation/inactivation of CaMKIIa-RNs on the efficacy of postural corrections stabilizing the trunk orientation, we calculated the coefficient of postural stabilization (Fig. 3**I**). We found that the mean value of the coefficient of stabilization during unilateral activation (Fig. 3**J**) as well as during unilateral inactivation (Fig. 3**K**) of CaMKIIa-RNs did not differ significantly from that in control. These results suggest that asymmetry in activity of CaMKIIa-RNs does not affect the efficacy of postural corrections.

### The roll tilt caused by asymmetry in activity of CaMKIIa-RNs is maintained during locomotion

Next, we addressed the question, whether the roll tilt caused by unilateral activation/inactivation of CaMKIIa-RNs is maintained during locomotion. As in other terrestrial quadrupeds (Zomlefer et al. 1984; Karayannidou et al., 2009; Misiaszek, 2006), in mice, there are left-right oscillations of the spine during locomotion with the maximal deviation of the spine toward the hindlimb that is in the beginning of the stance phase of locomotor cycle while the opposite hindlimb is at the end of the stance (at toe-off moment; Fig. 4**A**). To find out whether there is lateral displacement—as an indication of trunk roll tilt—of these spine oscillations during activation/inactivation of CaMKIIa-RNs as compared to control, first, we calculated the stabilized spine position before and after CNO administration (see Methods for details). Then the difference between the values of the stabilized spine positions observed after CNO administration and in control (the spine displacement) was calculated. We found that during unilateral activation of CaMKIIa-RNs, the spine displacement values were positive for most animals indicating that the spine was displaced toward the ipsilateral side and thus, ipsilateral body roll tilt was maintained. However, on average, the displacement was not significantly different from 0 (11±15%, *p* = 0.18, Fig. 4**D**). By contrast, during unilateral inactivation of CaMKIIa-RNs, the values of the spine displacement were negative, and the population average of the displacement was significantly different from 0 (-11±2%, *p* = 0.002; Fig. 4**D**). Thus, unilateral inactivation of CaMKIIa-RNs during locomotion evoked displacement of the spine toward the contralateral side suggesting that the trunk orientation with a contralateral roll tilt was maintained.

**Figure 4.**
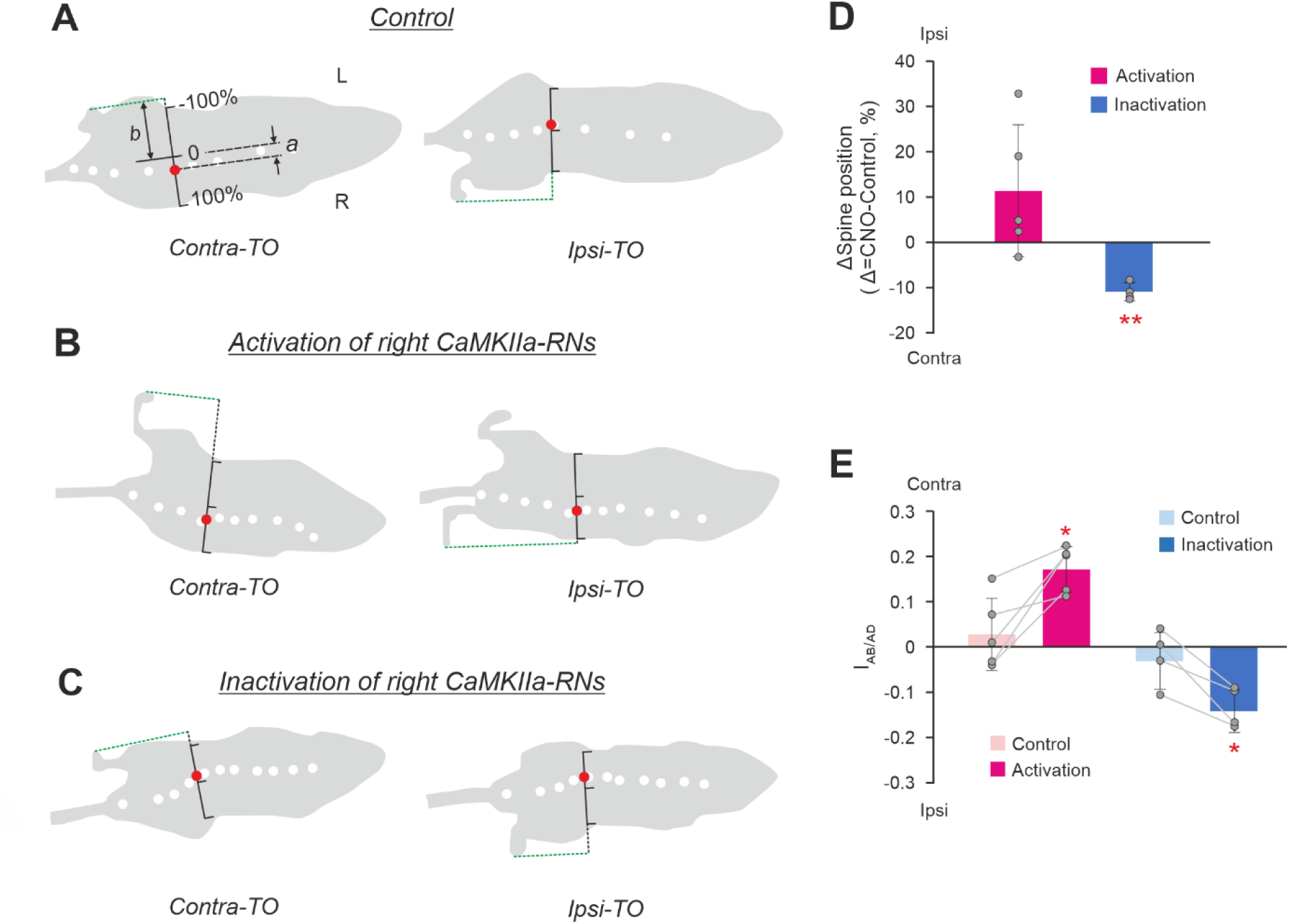
Effects of unilateral activation/inactivation of CaMKIIa-RNs on the body orientation during locomotion. **A,B,C.** Silhouettes of the top views of the walking mouse at the moments of the maximal displacement of the spine toward the ipsilateral hindlimb [at the moment of the contralateral hindlimb toe off (*Contra-TO*), left panels] and toward the contralateral limb [at the moment of the ipsilateral hindlimb toe off (*Ipsi-TO*), right panels] in control (**A**) and during activation (**B**) and inactivation (**C**) of the right CaMKIIa-RNs. White dots are markers on the spine. The red dot indicates the point on the spine that exhibits maximal left-right oscillations during locomotion. The black scale indicates the body width with its middle considered as “0” and the ipsilateral and contralateral edges of the body as +100% and -100%, respectively. **D.** The spine displacement (a difference of the spine position from the control value; see *Results* for details) during unilateral activation and inactivation of CaMKIIa-RNs. Values of the difference in individual animals, as well as the corresponding mean ± SD values are shown. **E.** Values of the abduction/adduction asymmetry index during locomotion (*I_A_B_/_AD*) in individual animals, as well as the corresponding mean ± SD values, before (*Control*) and during unilateral activation and inactivation of CaMKIIa-RNs. In **D,E**: *N* = 5 for unilateral activation and *N* = 4 for unilateral inactivation. In **D,E** Abbreviations as in Fig. 2.

To examine whether the asymmetry in the lateral position of the left and right limbs caused by the unilateral activation/inactivation of CaMKIIa-RNs in standing animals was maintained during locomotion, we calculated the abduction/adduction asymmetry index during locomotion in control and after activation/inactivation of CaMKIIa-RNs (see Methods for details). Positive and negative values of the index indicate displacement of the limbs from their symmetrical position in relation to the trunk toward the contralateral and ipsilateral side, respectively. We found that after unilateral activation of CaMKIIa-RNs, the values of the abduction/adduction asymmetry index were positive and the mean value of the index was significantly higher than that in control (respectively, 0.17±0.05 *vs* 0.03±0.08; paired *t* test, *p* = 0.02; Fig. 4**E**). These results suggest that unilateral activation of CaMKIIa-RNs evoked displacement of the hindlimbs toward the contralateral side in contrast to almost symmetrical limb position observed in control. On the other hand, after unilateral inactivation of CaMKIIA-RNs, values of the abduction/adduction asymmetry index were negative (Fig. 4**E**). On average, the mean value of the index differed significantly from that in control (respectively, -0.14±0.05 *vs* -0.03±0.06; paired *t* test, *p* = 0.02; Fig. 4**E**). These results suggest that unilateral inactivation of CaMKIIa-RNs evoked displacement of the hindlimbs toward the ipsilateral side as compared to control. Thus, during locomotion, lateral displacement of the limbs in relation to the trunk toward the contralateral side during CaMKIIa-RNs activation may contribute to maintenance of the trunk orientation with some roll tilt to the ipsilateral side, while displacement of limbs toward the ipsilateral side during CaMKIIa-RNs inactivation may contribute to maintenance of the contralateral tilt of the trunk.

### Asymmetry in activity of CaMKIIa-RNs hinders execution of righting behavior

The righting behavior (Magnus, 1924) that requires coordinated activity of left and right muscles of the trunk and limbs, is essential for control of posture. To find out whether asymmetry in CaMKIIa-RNs affects its execution, we compared the righting behavior before and during unilateral CaMKIIa-RNs activation/inactivation.

To evoke the righting behavior, we released animals in an upside-down position (moment 1 in Fig. 5**A**). In control, mice performed the righting in two Stages (Zelenin et al., 2021). During Stage 1, twisting and lateral bending (oblique bending) of the forequarters in relation to the hindquarters led to rotation of the body toward the side of twisting that was accompanied by movements of the forelimbs toward the surface. At the end of the Stage 1, the forequarters assumed a position with forelimbs standing on the surface, while the hindquarters turned from the upside-down position to the side (moment 2 in Fig. 5**A**). During Stage 2, the hindquarters rotated in relation to the forequarters until they reached a position close to the dorsal side up with hindlimbs standing on the surface (moment 3 in Fig. 5**A**).

**Figure 5.**
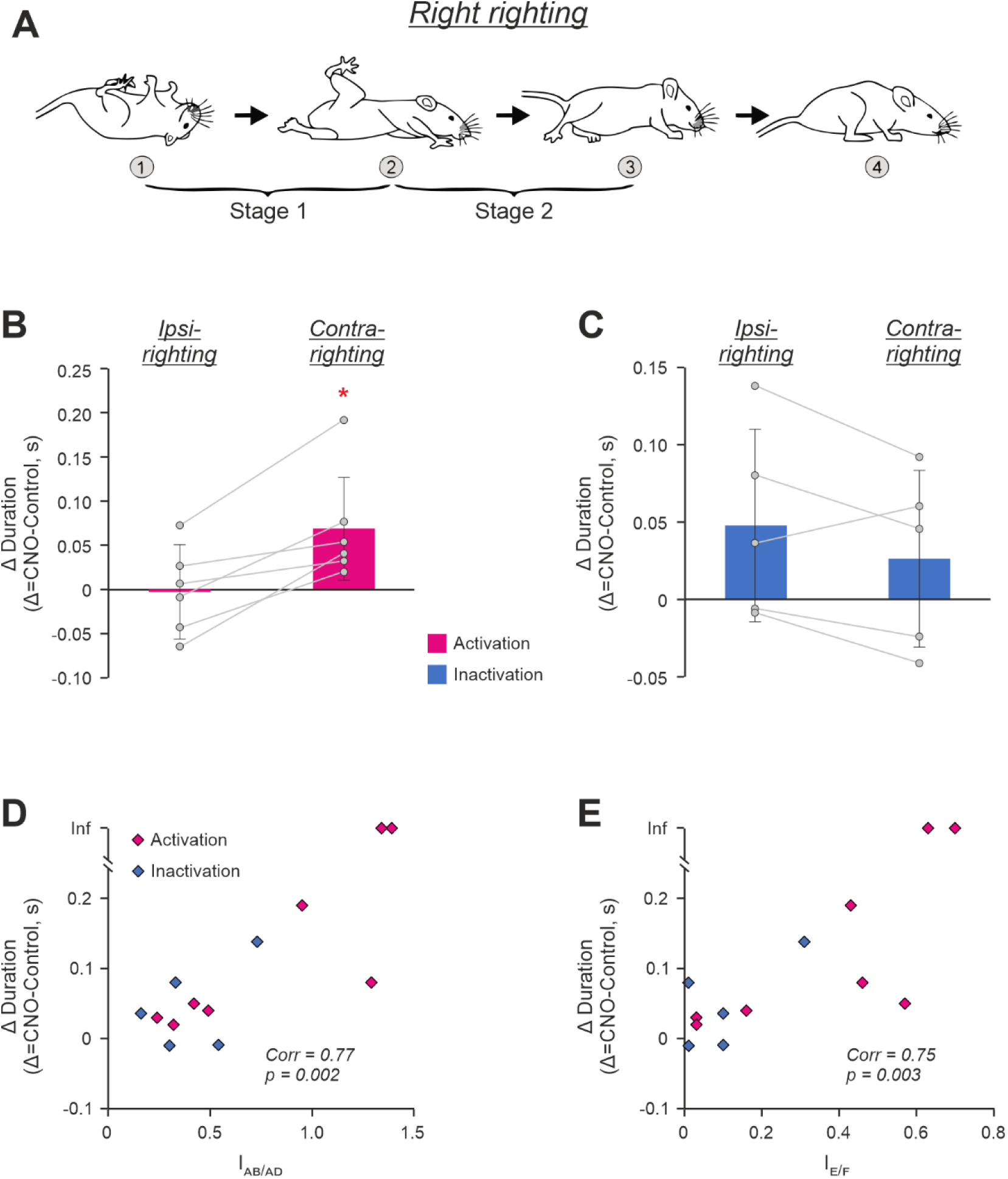
Effects of unilateral activation/inactivation of CaMKIIa-RNs on the righting behavior. **A.** Sequential positions of a mouse (*1-3*) that, starting from an upside-down position, acquired a dorsal side-up position. Two stages of the righting (*Stage 1* and *Stage 2*) are indicated. **B,C.** Differences between durations of the ipsilateral as well as contralateral righting (*Ipsi-* and *Contra-righting*, respectively) during unilateral activation (**B**) and inactivation (**C**) of CaMKIIa-RNs and the corresponding controls. Values of the difference in individual animals, as well as corresponding mean ± SD values are shown. **D.E.** Positive correlation between the abduction/adduction asymmetry index (*I_AB/AD_*in **D**) as well as the extension/flexion asymmetry index (*I_E/F_*in **E**) during unilateral activation/inactivation of CaMKIIa-RNs and the increase in duration of the righting performed toward the less active subpopulation of CaMKIIa-RNs as compared to control. *Corr*, Sperman’s rank correlation coefficient. Two animals that during unilateral activation of CaMKIIa-RNs were unable to perform the contralateral righting are indicated by points with the infinity (*Inf*) ordinate. In **B,C,D**, and E: N = 6, 5, 13, and 13, respectively.

Out of 13 tested mice, 11 mice were able to successfully perform both Stages of the righting with rotation to the ipsilateral (upward movement of the infected side) as well as to contralateral (downward movement of the infected side) side during both unilateral activation (N=6) and unilateral inactivation (N=5) of CaMKIIa-RNs. Two mice with excitatory DREADDs, during unilateral activation of CaMKIIa-RNs, performed ipsilateral righting but were unable to perform the righting to the contralateral side. We defined the righting duration as the sum of Stage 1 and Stage 2 durations.

We found that during unilateral activation of CaMKIIa-RNs, righting to the contralateral side was performed slower than in control. The mean value of difference in the righting duration during activation and in control (0.07±0.06 s) significantly differed from 0 (one-sample *t* test, *p* =0.045; Fig. 5**B**). However, the duration of the ipsilateral righting was similar to that in control (the difference in durations was -0.01±0.05 s, Fig. 5**B**). On the other hand, during unilateral inactivation of CaMKIIa-RNs, the difference in righting durations had positive values for both the ipsilateral and contralateral righting, indicating that they were slower than in control. However, difference in durations was slightly larger for ipsilateral righting than for the contralateral one, (respectively, 0.05±0.06 s and 0.03±0.06 s; Fig. 5**C**) suggesting that, in contrast to activation of CaMKIIa-RNs, righting to the ipsilateral side was affected stronger than that to the contralateral side.

Thus, there was a tendency that left-right asymmetry in activity of CaMKIIa-RNs led to an increase in the duration of the righting performed toward the less active subpopulation of CaMKIIa-RNs. To clarify whether this increase in the duration was caused by asymmetry in the left-right limb configurations, we plotted the abduction/adduction asymmetry index as well as the extension/flexion asymmetry index against the change in the righting duration performed toward the side with lower CaMKIIa-RN activity (Fig. 5, **D** and **E**, respectively). We found a significant positive correlation between parameters in both cases suggesting that asymmetry in configurations of the left and right limbs caused by unilateral activation/inactivation of CaMKIIa-RNs – extension and abduction of the hindlimb on the side of the less active subpopulation of CaMKIIa-RNs and simultaneous flexion and adduction of the hindlimb on the opposite side – distorted righting reflex toward the side with lower CaMKIIa-RN activity.

### The body roll tilt was caused specifically by CaMKIIa-RNs located in MdV-IRt area of the caudal medulla

To clarify whether the effects of CaMKIIa-RNs on the body posture were specific for MdV– IRt area of the caudal medulla, we studied effects of unilateral activation of CaMKIIa-RNs in the gigantocellular reticular nuclei, that is located rostrally to MdV-IRt (Gi, Fig. 6**A**) and in the area that is dorsal to MdV-IRt (d-MdV, Fig. 6**D**). Figure **6B,C****,E,F** show representative examples of areas infected in Gi (**B**) and in d-MdV (**E**), as well as the average infected areas in Gi (**C**) and in d-MdV (**F**) presented as heatmaps.

**Figure 6.**
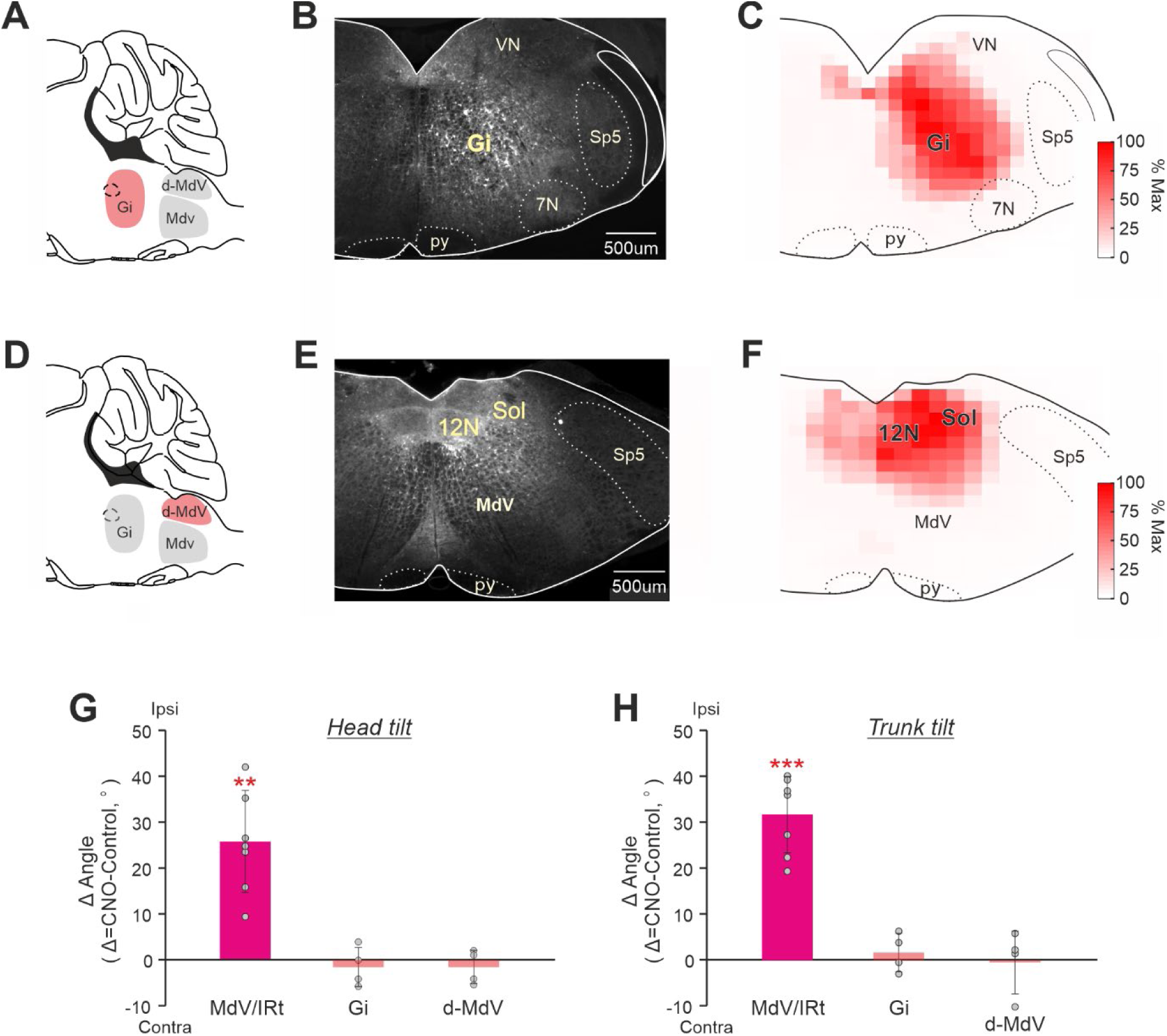
Comparison of effects of unilateral activation of CaMKIIa-RNs located in MdV-IRt area and in two adjacent areas. **A,D.** A scheme of the sagittal section of the brainstem indicating the location of two areas (the gigantocellular nucleus, Gi, and the area that is dorsal to MdV, d-MdV) adjacent to MdV-IRt area. **B,E.** Representative examples of unilateral infection of CaMKIIa-RNs in Gi (**B**) and in d-MdV (**E**) with AAV-CaMKIIa-hM3Dq-mCherry. **C,F.** Heatmaps showing the averaged extent of the infected area in Gi (**C**, *N* = 4) and in d-MdV (**F**, *N* = 4). **G-H.** Comparison of effects of unilateral activation of CaMKIIa-RNs in MdV-IRt, in d-MdV, and in Gi area on the head (**G**) and trunk (**H**) roll tilt in animals standing on a horizontal surface. In **G** and **H**, mean ± SD values of the difference between two conditions (unilateral activation and control) are shown. In **G,H:** *N* = 7 for MdV-IRt, *N* = 4 for Gi, and *N* = 4 for d-MdV.

We found that unilateral activation of CaMKIIa-RNs in these two areas produced no significant effects on the body posture. During standing on a horizontal surface, for the roll tilt angles of the head and trunk, difference from the corresponding control values was on average close to zero (Fig. **6G,H**). These results differed dramatically from those obtained during unilateral activation of CaMKIIa-RNs in MdV–IRt area (Fig. 6**G,H**), suggesting that CaMKIIa-RNs in MdV–IRt area but not in the adjacent areas, contribute to control of body orientation in the transverse plane.

Next, we addressed the question whether the effects on the body posture are specific to the molecular identity of CaMKIIa-RNs in MdV-IRt area. To answer this question, first, we compared the effects of unilateral activation of a broad population of glutamatergic (Vglut2) neurons and CaMKIIa-RNs in MdV-IRt area. For this purpose in Vglut2-Cre mice we infected Vglut2 reticular neurons (Vglut2-RNs) in MdV-IRt area with AVV-hsyn-lox-hM3Dq-mCherry-lox (Fig. 7**A**,**B**). Figure 7**D**-**H** compares effects of unilateral activation of Vglut2-RNs and CaMKIIa-RNs in MdV-IRt area. While activation of CaMKIIa-RNs evoked the ipsilateral roll tilt of the body (reflected in displacement of the spine toward the ipsilateral side of the body; right panels in Fig. 7**G,H**), activation of Vglut2-RNs caused the ipsilateral bending of the body in the yaw plane (left panels in Fig. 7**G**,**H**). Also, during unilateral activation of CaMKIIa-RNs, the animal was able to stand still without the head or limb movements (Fig. 7**D,E**, lower panels). By contrast, during unilateral activation of Vglut2-RNs, continuous movements of the head and ipsilateral forelimb (Fig. 7**D**,**E**, upper panels) accompanied standing. Finally, during unilateral activation of Vglut2-RNs, the animals performed continuous ipsilateral turning (circling) during locomotion in the open field (Fig. 7**F**, upper panel), while during unilateral activation of CaMKIIa-RNs, animals performed locomotion with right and left turns which randomly occurred in approximately equal proportion (Fig. 7**F**, lower panel). These differences in effects of unilateral activation of CaMKIIa-RNs and Vglut2-RNs were observed in all studied animals (*N*=7 for CaMKIIa-RNs and *N*=3 for Vglut2-RNs).

**Figure 7.**
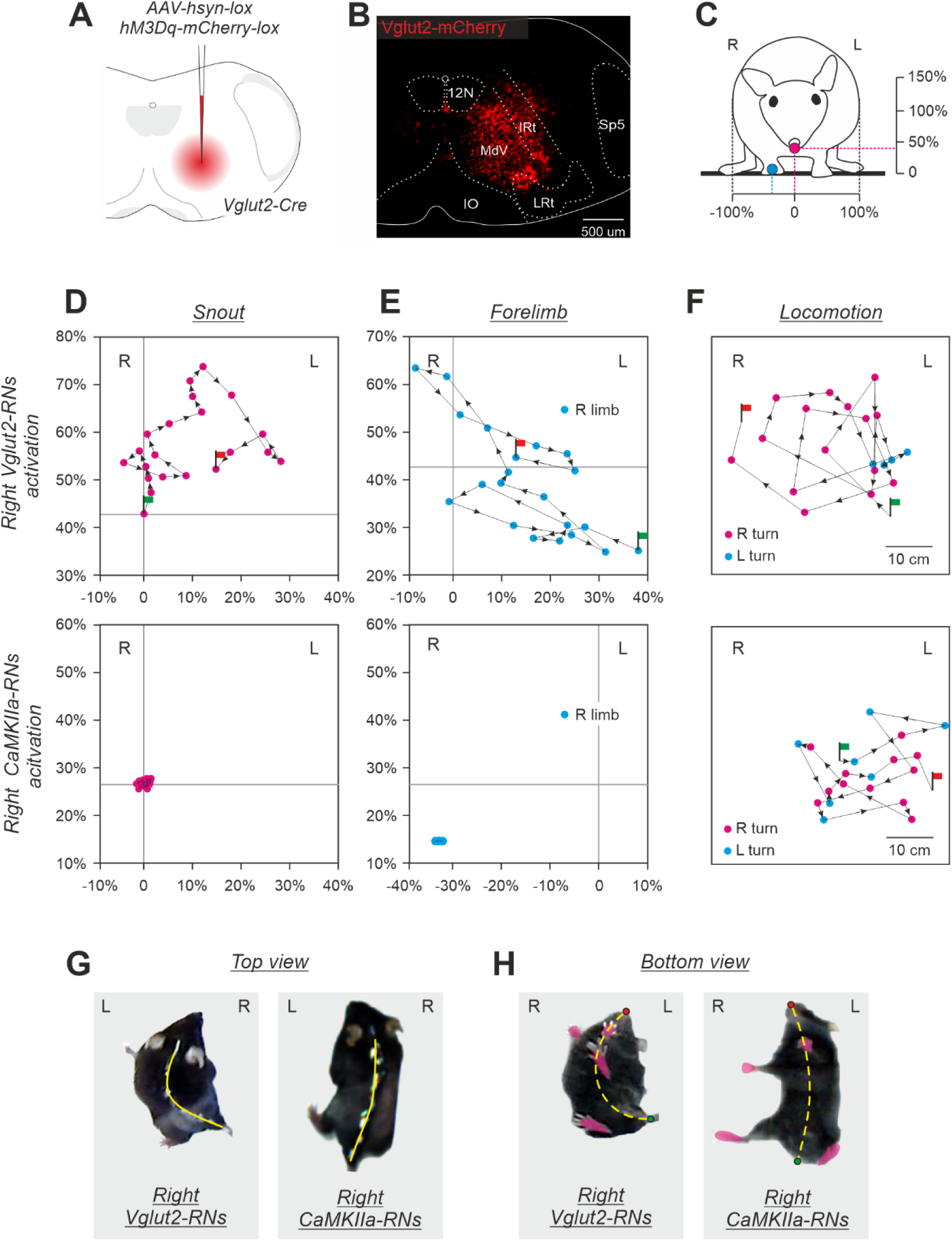
Comparison of effects of unilateral activation of CaMKIIa-RNs and Vglut2-RNs located in MdV-IRt area. **A,B.** Unilateral infection of Vglut2-RNs with AAV-hsyn-lox-hM3Dq-mCherry-lox in the Vglut2-Cre mouse. **C.** The front view of a mouse. Crimson and cyan circles indicate the positions of the snout and ipsilateral forelimb, respectively. The horizontal and vertical axes for positions of the snout (**D**) and the forelimb (**E**) are shown. The right and left edges of the body correspond, respectively, to -100% and +100% of the half of the body width; 0 for the vertical axis is at the support surface; the vertical scale is the same as the horizontal one. **D**,**E**. Comparison of trajectories of the snout (**D**) and the right forelimb (**E**) movements in the transverse plane in the animal with unilateral activation of the right Vglut2-RNs (upper panels) and in the animals with unilateral activation of the right CaMKIIa-RNs (lower panels); 0,02 s between points. **F**. Comparison of trajectories of locomotion in the open field performed by the mouse during activation of the right Vglut2-RNs and by the mouse during activation of the right CaMKIIa-RNs; 1 s between points. In **C-F,** Green and red flag indicate the beginning and the end of the trajectory, respectively. **G**,**H**. Top and bottom views of the mouse during activation of the right Vglut2-RNs and mouse during activation of the right CaMKIIa-RNs. In **G** and **H**, the position of the spine and the midline of the body are shown by solid and dashed yellow lines, respectively.

Second, we compared the effects of unilateral activation of inhibitory GABA-ergic neurons and CaMKIIa-RNs in MdV-IRt area. For this purpose, in GAD67-Cre mice we infected GAD67 reticular neurons (GAD67-RNs) in MdV-IRt area with AVV-hsyn-lox-hM3Dq-mCherry-lox (Fig. 8**A**,**B**). Figure 8**C**,**D** compares effects of unilateral activation of GAD67-RNs and CaMKIIa-RNs in MdV-IRt area. While the differences between head and trunk roll tilt angles during unilateral activation of CaMKIIa-RNs and control were significant, they were insignificant during unilateral activation of the GAD67-RNs (respectively, –4±5° and –1±4°, one-sample *t* test, *p* = 0.09, and *p* = 0.71; Fig. 8**C**,**D**). Thus, left-right asymmetry in activity of GABA-ergic neurons in MdV-IRt area does not affect the body orientation in the transverse plane.

**Figure 8.**
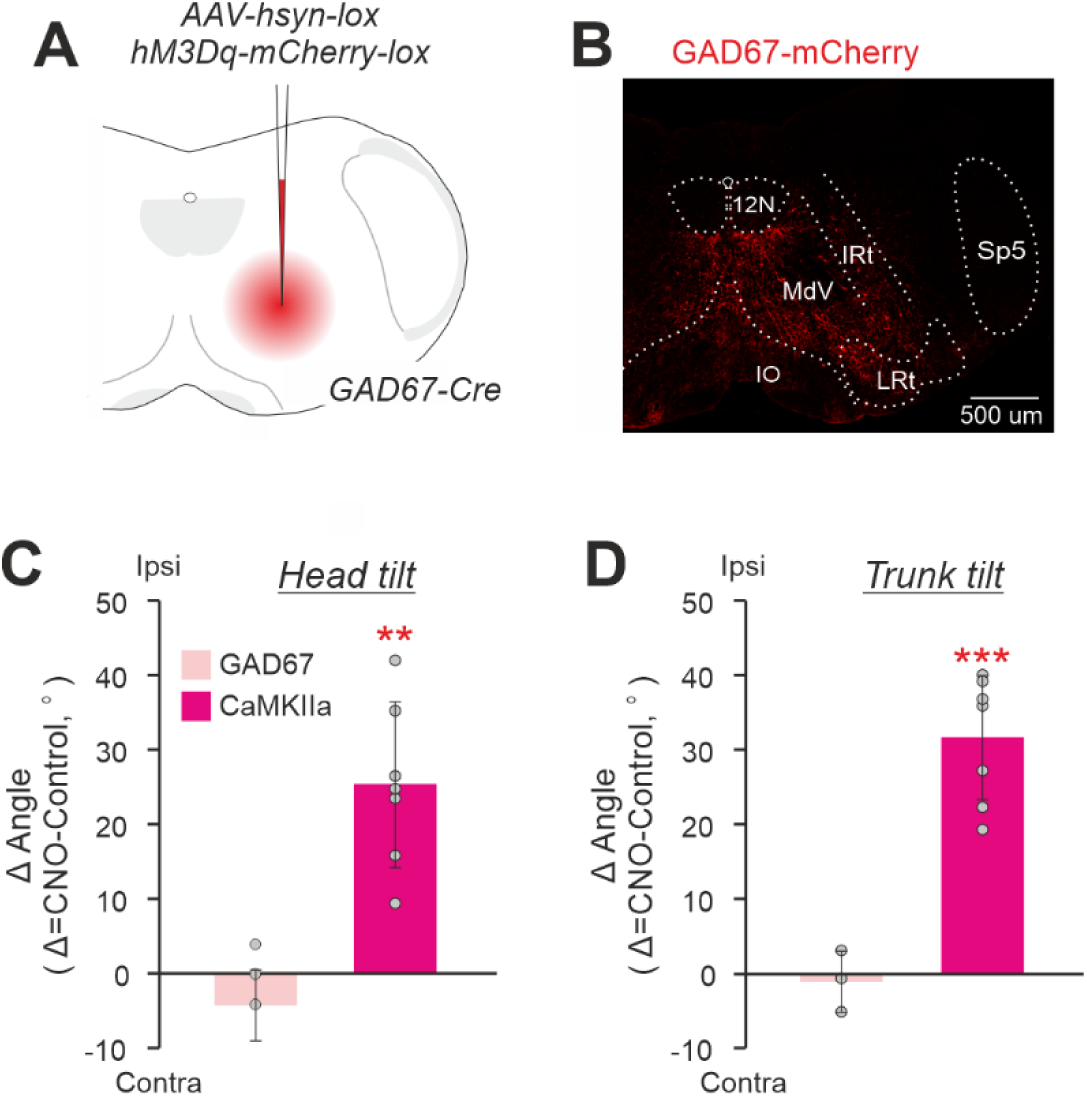
Comparison of effects of unilateral activation of CaMKIIa-RNs and GAD67-RNs located in MdV-IRt area. **A,B.** Unilateral infection of GAD67-RNs with AAV-hsyn-lox-hM3Dq-mCherry-lox in the GAD67-Cre mouse.**C,D.** Comparison of effects on the head (**C**) and trunk (**D**) roll tilt caused by unilateral activation of GAD67-RNs. Values of averaged difference between the roll tilt angle observed during unilateral activation of neurons and in control (*Δ Angle*) are shown for individual animals as well as corresponding mean ± SD values. In **C,D**: *N* = 3 for GAD67-RNs and *N* = 7 for CaMKIIa-RNs.

Taken together, all these results suggest that the change of the body orientation in the transverse plane (the body roll tilt) is caused by left-right asymmetry in activity of a specific molecularly identified population of RNs located in a definite brainstem area. It is a population of RNs expressing CaMKIIa and located specifically in MdV-IRt area.

### The majority of CaMKIIa-RNs in MdV-IRt area are glutamatergic

To reveal neurotransmitter phenotypes of CaMKIIa-RNs located in MdV-IRt area, first, AAV-CaMKIIa-GFP viruses were injected in MdV-IRt area to label CaMKIIa-RNs with GFP (upper panels in Fig. 9**A**,**B**). Then by using RNAscope *in situ* hybridization for vesicular glutamate transporter 2 (Vglut2) and vesicular inhibitory amino acid transporter (Vgat), Vglut2 positive and Vgat positive neurons in MdV-IRt area were identified (middle panels in Fig. 9**A**,**B**). We found both neurons with co-expression of GFP and *Vglut2* (lower panel in Fig. 9**A**), as well as neurons with co-expression of GFP and *Vgat* (lower panel in Fig. 9**B**) in MdV-IRt area. Thus, the population of CaMKIIa-RNs contains both excitatory glutamatergic and inhibitory GABAergic/glycinergic neurons. However, the relative number of excitatory CaMKIIa-RNs was significantly (more than twofold) higher than the relative number of the inhibitory ones (respectively, 64±11% *vs* 26±11%, unpaired *t* test, *p* = 7×10^-4^; Fig. 9**E**). The glutamatergic and inhibitory CaMKIIa-RNs were intermixed (Fig. 9**C**,**D**) and rather similarly distributed in MdV-IRt area (Fig. 9**F**,**G**).

**Figure 9.**
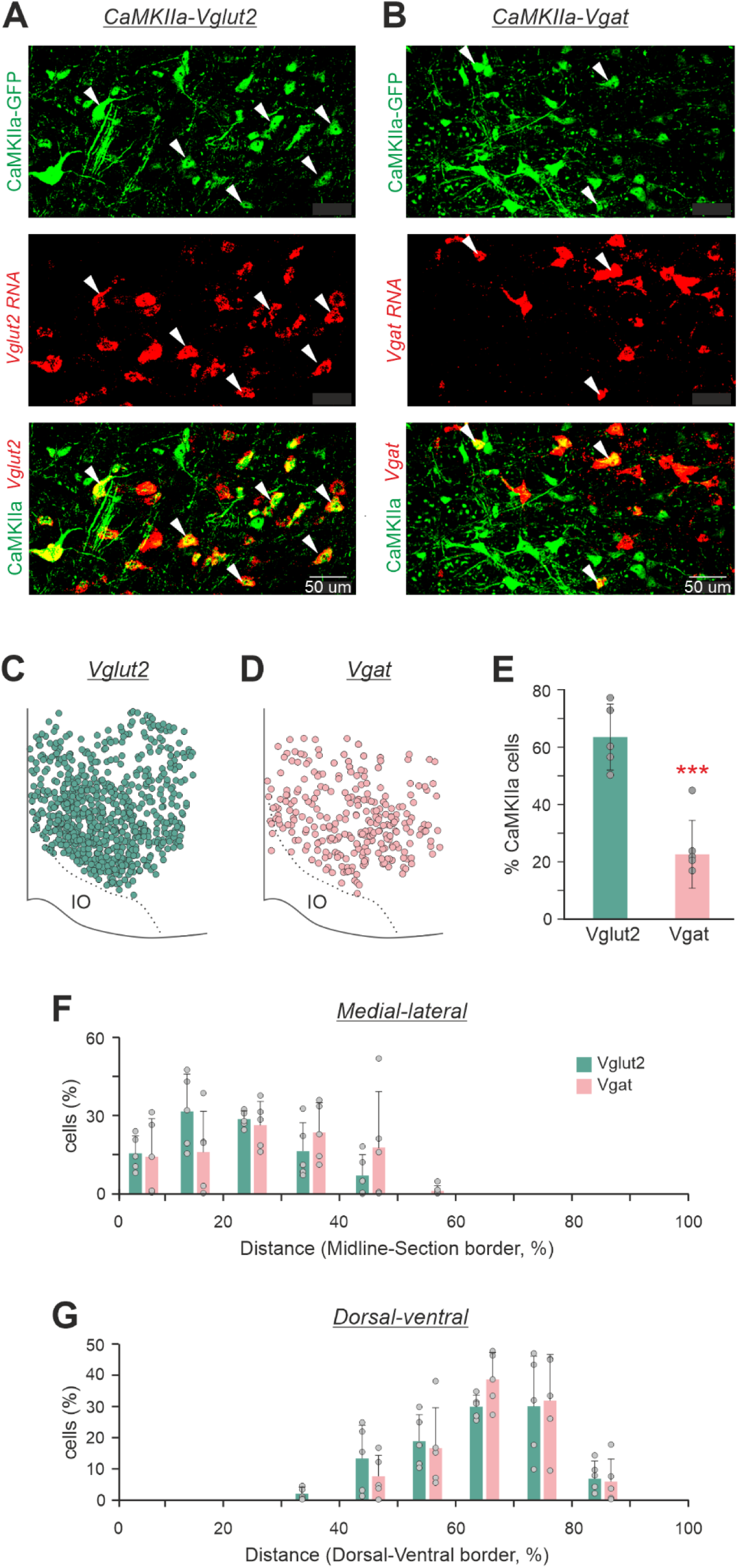
Neurotransmitter phenotypes of CaMKIIa-RNs located in MdV-IRt area. **A,B.** Two representative images with GFP expression in CaMKIIa-RNs (green cells in the upper panels), with *in situ* hybridization for Vglut2 mRNA identifying glutamatergic RNs (**A**, red cells in the middle panel), with *in situ* hybridization for Vgat mRNA identifying GABAergic RNs (**B**, red cells in the middle panel), with co-expression of GFP and for Vglut2 mRNA identifying glutamatergic CaMKIIa-RNs (**A**, yellow cells in the lower panel), and with co-expression of GFP and for Vgat mRNA identifying GABAergic CaMKIIa-RNs (**B**, yellow cells in the lower panel). White arrowheads in **A** and **B** mark glutamatergic and GABAergic CaMKIIa-RNs, respectively. **C,D.** Location of glutamatergic (**C**) and GABAergic (**D**) CaMKIIa-RNs in MdV-IRt area. **E.** Percentage of glutamatergic (Vglut2) and GABAergic (Vgat) CaMKIIa-RNs in MdV-IRt area. **F,G.** Percentage of Vglut2^+^ and percentage of Vgat^+^ CaMKIIa-RNs in MdV-IRt area at different medio-lateral (**F**) and dorso-ventral (**G**) levels. For identification of Vglut2 CaMKIIa-RNs and Vgat CaMKIIa-RNs: *N* = 3, *n* = 5 sections, 683 neurons, and *N* = 3, *n* = 5 sections, 254 neurons, respectively.

### Population of CaMKIIa-RNs located in MdV-IRt area contains reticulospinal neurons

To determine whether the population of CaMKIIa-RNs in MdV-IRt area contains reticulospinal neurons, we examined presence of the mCherry signals in the spinal cord sections of mice with unilateral injection of AAV-CaMKIIa-hM3Dq-mCherry in MdV-IRt area. We found mCherry^+^ axons at the cervical, thoracic and lumbar levels of the spinal cord (Fig. 10**A**-**C**,**D**-**F**), demonstrating that the population of CaMKIIa-RNs in MdV-IRt area contains reticulospinal neurons. In the spinal cord, mCherry^+^ axons descended mainly in the medial part of the ipsilateral lateral funiculus and their number decreased from cervical to lumbar level (Fig. 10**A**-**F**). At all three levels of the spinal cord, the greatest arborization of mCherry^+^ axons was found within the intermediate part of the ipsilateral gray matter in laminae VII, VIII and X (Fig. 10**G**-**I**). Also, at the cervical and thoracic levels a substantial arborization was found in the same laminae of the contralateral gray matter (Fig. 10**A**,**B**,**G**,**H**). Notably, mCherry^+^ terminals were largely absent in the dorsal horn laminae I–VI where sensory networks are localized, and in lamina IX where the majority of limb muscle motoneurons reside. Since lamine VII and VIII contain interneurons and laminae X contains motoneurons of the axial muscles, one can conclude that CaMKIIa-reticulospinal neurons most likely directly affect predominantly interneurons as well as motoneurons of the axial muscles at all levels of the spinal cord.

**Figure 10.**
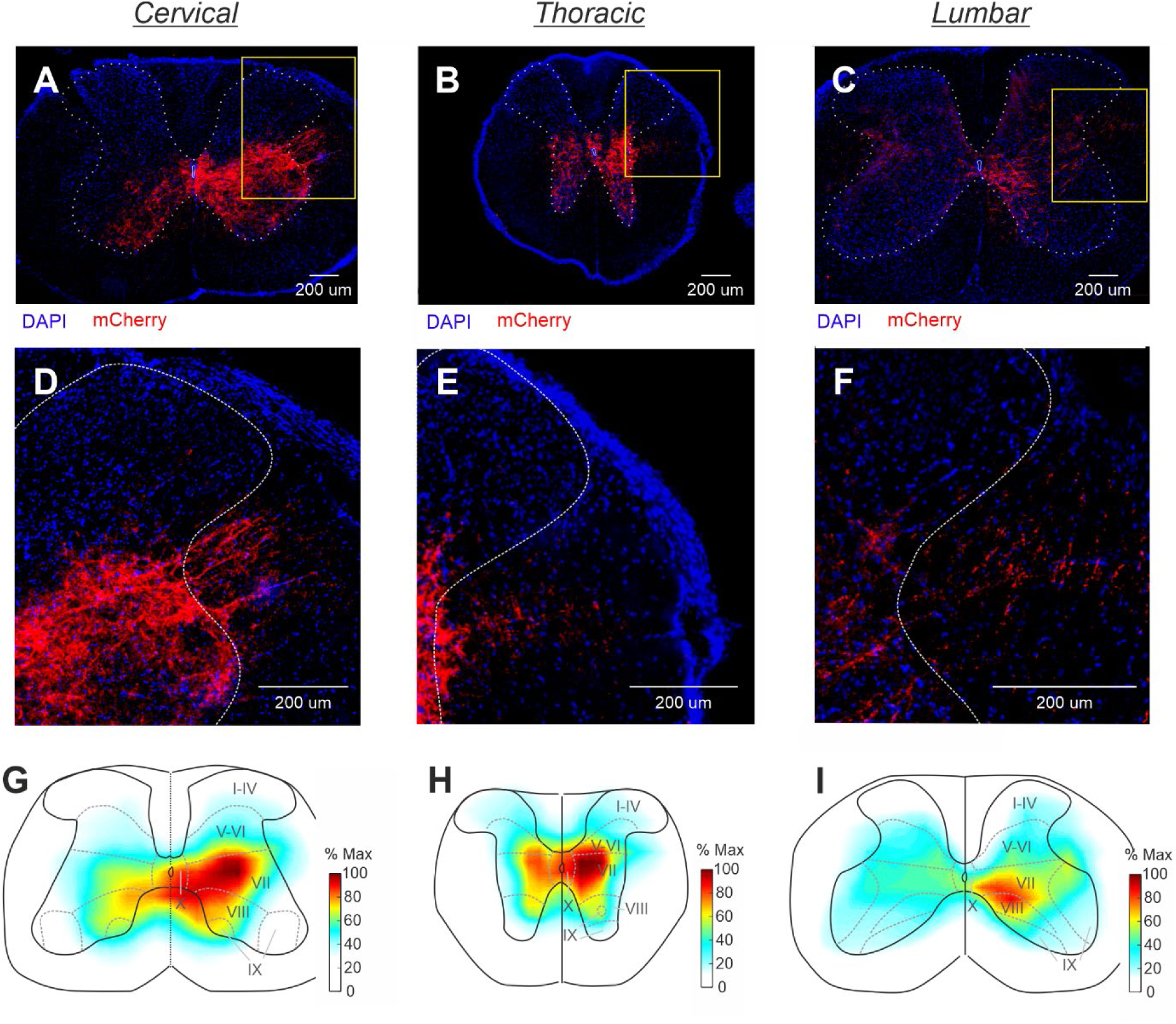
Population of CaMKIIa-RNs in MdV-IRt area contains reticulospinal neurons. **A-F.** A representative example of mCherry^+^ axons located mainly in the medial part of the ipsilateral lateral funiculus, as well as their arborizations in the gray matter at the cervical (**A,D**), thoracic (**B,E**), and lumbar (**C,F**) levels of the spinal cord. Yellow rectangles in **A-C** delineate the areas shown in **D-F** with larger amplification. **G-I.** A heatmap showing averaged fluorescent intensity (% Max) of mCherry^+^ axons and their arborizations in the cervical (**G**), thoracic (**H**), and lumbar (**I**) spinal cord (*N* = 2, *n* = 10, 12, and 18, for cervical, thoracic, and lumbar spinal cord, respectively).

## DISCUSSION

In the present study we identified a population of reticular neurons located in the caudal medulla – CaMKIIa-RNs in MdV-IRt area – which controls the body orientation maintained by an animal in the transverse plane.

Although it was reported that the caudal medulla contains rather sparse CaMKIIa neurons (Wang et al., 2013), we demonstrated that unilateral chemogenetic activation or inactivation of the CaMKIIa-RNs located MdV-IRt area elicited a lateral body sway towards the dominant (more active) side, and this new body orientation was actively stabilized during standing and maintained during locomotion. Earlier, it was suggested that terrestrial quadrupeds stabilize the body orientation in the transverse plane due to continuous interaction of antagonistic postural limb reflexes (PLRs) generated by the left and right limbs (Beloozerova et al., 2003; Hsu et al., 2012; Fig. 11). The animal stabilizes such orientation at which the effects of antagonistic PLRs are equal to each other. This is a set-point of the postural control system —the dorsal side-up body orientation that the animal actively stabilizes. Any deviation from this orientation leads to enhancement of PLRs generated by the left or right limbs, which return the body to the initial orientation (Fig. 11**A**,**B**). It was also demonstrated that left-right asymmetry in the tonic vestibular input caused by binaural galvanic vestibular stimulation (GVS) that presumably creates left-right asymmetry in the vestibulospinal and reticulospinal systems, evokes a shift of the set-point of the postural system through a change of gains in antagonistic PLRs (Fig. 11**C**,**D**). This leads to stabilization of a new orientation of the body with some roll tilt to the anode (Beloozerova et al., 2003; Hsu et al., 2012). Results of the present study support the hypothesis about the role of the reticulospinal system in control of the body orientation in terrestrial quadrupeds. We suggest that left-right asymmetry in activity of the population of CaMKIIa-RNs located MdV-IRt area, which contains reticulospinal neurons, shifts the set-point of the antagonistic PLRs leading to stabilization of a new body orientation with some roll tilt towards the dominant side (Fig. 11**C,D**). In the natural environment, such shift of the set-point of the postural system allows the animal to stabilize the dorsal side-up body orientation on a laterally inclined surface (Fig. 11**F**). A similar principle of balance control was also found in simpler animals – a mollusk (Clione) and a lower vertebrate (lamprey) (Deliagina et al., 1998; Deliagina and Fagerstedt, 2000; Deliagina et al., 2014). In the lamprey, antagonistic postural reflexes are mediated by two populations of reticulopsinal neurons, and different factors that produce asymmetry in their activity, affect the body orientation stabilized in the transverse plane.

**Figure 11.**
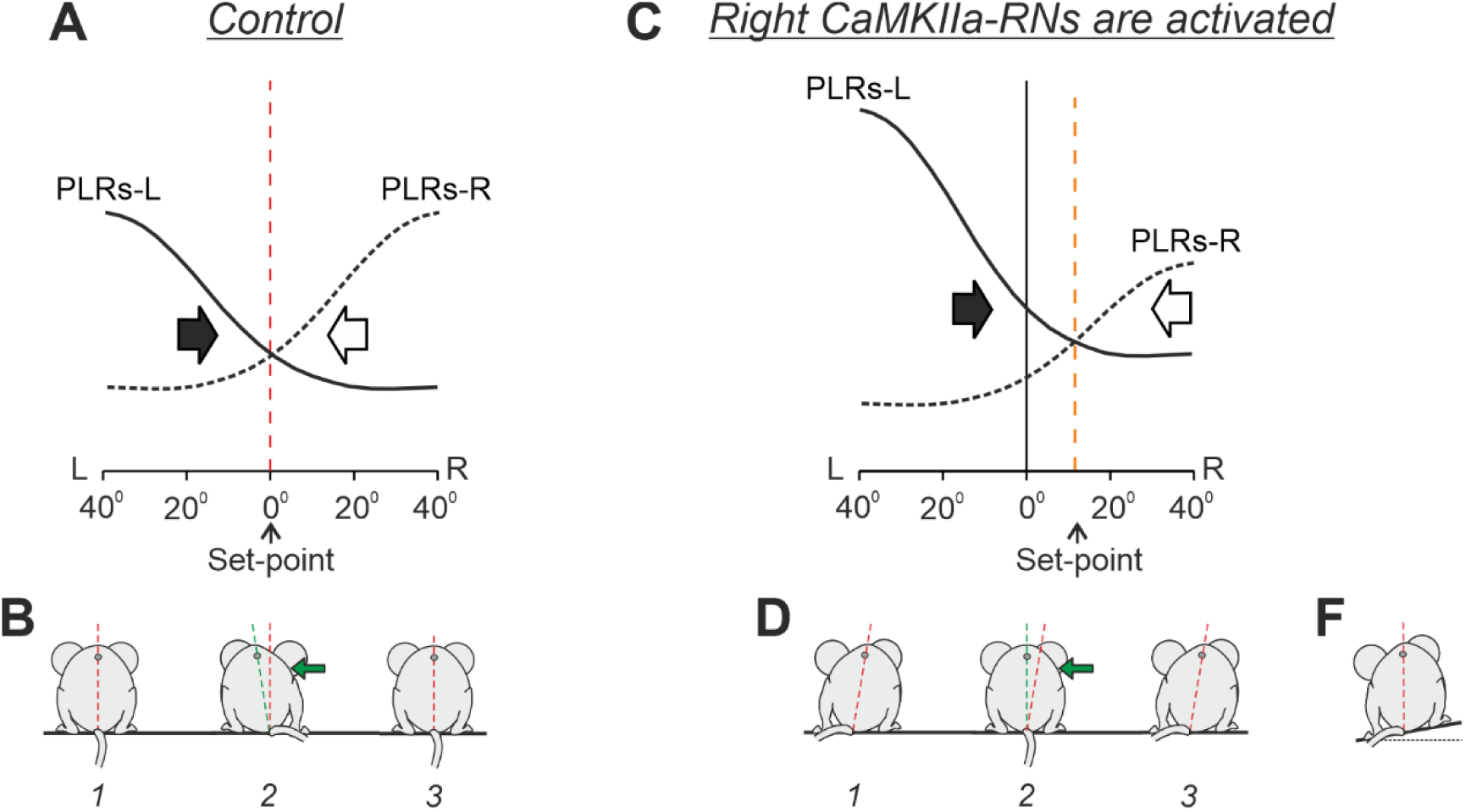
Conceptual model of the trunk stabilization system and effects of left-right asymmetry in activity of CaMKIIa-RNs. **A-D.** Presumed effects of the two antagonistic postural limb reflexes (PLRs) in the unrestrained standing mouse in control (**A** and **B**) and during activation of right CaMKIIa-RNs (**C** and **D**). **A** and **C**: the abscissa shows a deviation of the dorso-ventral body axis from the vertical (lateral sway); the ordinate shows the values of the right and left (L) PLRs (PLR-R and PLR-L, solid and dashed lines, respectively). Black and white arrows indicate the motor effect (lateral sway) caused by PLR-R and PLR-L, respectively. **B** and **D**. The stabilized orientation (*1*), effect of the lateral push (*2*), and the restored orientation (*3)*. **F.** The shift of the set- point of the postural control system caused by the left-right asymmetry in activity of CaMKIIa-RNs allows to stabilize the dorsal side-up orientation on the laterally inclined surface. (See DISCUSSION for details).

The fact that not only unilateral activation but also unilateral inactivation of CaMKIIa-RNs creates asymmetry in their activity sufficient for the behavioral effects suggests that the population of CaMKIIa-RNs in MdV-IRt area has a substantial level of activity in standing animals. This finding supports our suggestion about their important role in control of the body orientation. However, unilateral inactivation of CaMKIIa-RNs in standing animals caused much weaker behavioral effects as compared to those caused by unilateral activation. In particular, it was reflected in much smaller absolute values of the head and trunk roll tilts. Also, unilateral inactivation does not affect significantly the duration of the contralateral righting, while unilateral activation caused a significant increase in its duration. One can suggest that during quiet standing or during righting, the CaMKIIα-MdV neurons have a relatively low level of activity and thus their unilateral activation and inactivation create, respectively, a strong and weak left-right asymmetry in their activity leading, consequently, to strong and weak behavioral effects.

By contrast, during locomotion, unilateral inactivation of CaMKIIa-RNs evoked a significant body roll tilt, while the effect of unilateral activation on average was insignificant (although strong in some individual animals). One can suggest that during locomotion, CaMKIIa-RNs are strongly activated. Thus, their unilateral inhibition creates a strong left-right asymmetry in the activity of the population that leads to execution of a significant behavioral effect. By contrast, their unilateral activation creates a very weak (if any) asymmetry in the population activity (since the neuronal activity is already close to its maximal level) which results in absence of the behavioral effect.

We showed that the body roll tilt caused by unilateral activation/inactivation of CaMKIIa-RNs located MdV-IRt area is evoked due to flexion and adduction of the limbs on the dominant side and simultaneous extension and abduction of the opposite limbs. These results are in line with results of the earlier study demonstrated that electrical microstimulation of specific sites in the pontomedullar reticular formation evoked inactivation and activation of extensors in the ipsilateral and contralateral hindlimb, respectively (Takikusaki et al., 2016).

We demonstrated that the behavioral effect (the body roll tilt) evoked by the left-right asymmetry in activity of CaMKIIa-RNs is specific to the population of CaMKIIa-RNs located in MdV-IRt area, but not in the neighboring areas. Furthermore, within MdV-IRt area, this behavioral effect is evoked by specifically CaMKIIa-RNs, but not by the broader glutamatergic (Vglut2-RNs) or GABAergic (GAD67-RNs) populations. We found that the left-right asymmetry in activity of Vglut2-RNs located in MdV-IRt area evoked another behavioral effect that is the body bend in the yaw plane toward the dominant side accompanied by continuous movements of the head and the ipsilateral forelimb. Earlier, it was demonstrated that ablation of Vglut2-RNs located in MdV distorts skilled forelimb motor tasks (Esposito et al., 2014) and also affects orofacial movements (Kleinfeld et al. 2023). It was demonstrated that both the Vglut2 and inhibitory neurons in IRt contribute to rhythmic orofacial motor behaviors, such as whisking and licking (Kleinfeld et al., 2023). Thus, MdV-IRt area of the caudal medulla contains a number of different molecularly identified populations of RNs that control specific aspects of the motor behavior in mice.

We found that the majority of neurons in the population of CaMKIIa-RNs in MdV-IRt area are excitatory glutamatergic (Vglut2) neurons although it also contains inhibitory GABAergic/glycinergic (Vgat) neurons. These results are in line with results of the previous studies that documented that MdV contains both glutamatergic and inhibitory neurons (Esposito et al., 2014), and that CaMKIIa neurons can be excitatory as well as inhibitory (Wang et al., 2013). A goal of future studies will be to elucidate whether the evoked roll tilt of the body is due to left-right asymmetry in activity of only excitatory CaMKIIa-RNs, or only inhibitory CaMKIIa-RNs, or both excitatory and inhibitory CaMKIIa-RNs.

We demonstrated that the population of CaMKIIa-RNs located in MdV-IRt contains reticulospinal neurons with axons descending ipsilaterally and branching at all levels of the spinal cord with the intensity of branching in the cervical region higher than in lumbar region. Reticulospinal neurons originating from MdV with similar projections were described earlier (Esposito et al., 2014; Xie et al., 2023). However, while reticulospinal neurons originating from MdV formed abundant synaptic connections with limb motorneurons (Esposito et al., 2014; Xie et al., 2023), terminals of the population of CaMKIIa reticulospinal neurons originating from MdV-IRt area avoid lamina IX where the majority of limb motoneurons reside, suggesting that their effects on limb motoneurons are mediated by spinal interneurons. A recent study demonstrated a significant input from the MdV neurons to spinal V1 interneurons, further supporting the suggestion that projections from this region target spinal interneurons (Chapman et al. 2024). A question for future studies is whether the body roll tilt is evoked by the left-right asymmetry in activity of only CaMKIIa reticulospinal neurons, or only CaMKIIa-RNs that do not project to the spinal cord, or the entire population of CaMKIIa-RNs.

It is well documented that at the acute stage of incomplete spinal cord injury the motor functions (including postural functions) are severely distorted or absent. However, they gradually recover over time due to plastic changes in the corresponding neuronal networks (Hultborn & Malmsten, 1983a,b; Helgren & Goldberger, 1993; Kuhtz-Buschbeck *et al*. 1996; Lyalka et al., 2005; Frigon & Rossignol, 2006; Zelenin et al., 2023). It was demonstrated that reticulospinal neurons contribute to these plastic changes (Zörner et al., 2014; Asboth et al., 2018; Engmann et al., 2020). Since calcium–calmodulin-dependent protein kinase IIa is central to synaptic plasticity (Hinds et al., 2003; Lisman et al., 2012), one can expect that CaMKIIa-RNs from MdV-IRt area may contribute to plastic changes underlying the recovery of postural functions after the spinal cord injury.

To conclude, in the present study, for the first time, a population of CaMKIIa-RNs in MdV-IRt area was characterized and its functional role was demonstrated. We found that left-right asymmetry in activity of this population evoked the body roll tilt which was actively stabilized during standing and maintained during locomotion. We suggest that CaMKIIa-RNs in MdV-IRt area control the body orientation in the transverse plane. To maintain the dorsal-side-up body orientation during standing on a horizontal surface, the activity level of the right and left CaMKIIa-RNs must be equal to each other. On the other hand, to maintain the dorsal-side-up orientation on a laterally inclined surface, a right/left asymmetry in the CaMKIIa-RNs activity is necessary. We found that most CaMKIIa-RNs are excitatory, and that the population contains reticulospinal neurons. Obtained results advance our understanding of the neuronal mechanisms underlying stabilization of the body orientation at different environmental conditions.

## METHODS

### Animals

Experiments were performed on wild type (C57Bl6, *N*=24) mice, as well as on Vglut2-Cre (*N*=3) and GAD67-Cre (*N*=3) transgenic mice. The mice were of both sex, 18-30 g weight, and 8-12 weeks old. They were housed in standard cages with food and water *ad libitum* at a 12h light/12h dark cycle. All experiments were conducted with approval of the local ethical committee (Norra Djurförsöksetiska Nämnden) in Stockholm and followed the European Community Council Directive (2010/63EU) and the guidelines of the National Institute of Health Guide for the Care and Use of Laboratory animals.

### Stereotaxic viral injections

All surgical procedures were performed under general anesthesia and aseptic conditions. General anesthesia consisted of ketamine (75 mg/kg) in combination with medetomidine (1 mg/kg) administered intraperitoneally. The level of anesthesia was controlled by applying pressure to a paw (to detect limb withdrawal), and by examining the size and reactivity of pupils. Anesthetized mice were fixed in a stereotaxic frame. Mice were kept on a 37°C heating pad for the duration of the surgery. Viscotears were applied to the eyes to prevent dehydration. The skin on the head was shaved, and an incision was made to expose the skull. Skull references were taken for bregma and lambda. A small hole was drilled in the skull overlying the target brain region for injection. The underlying dura was opened. A pulled glass capillary was filled with mineral oil, secured to a capillary nanoinjector that was fixed on a micromanipulator. Viruses were mixed with a small amount of Fast Blue for visualization and loaded into the capillary. The capillary was advanced to the target brain region at a rate of 0.1 mm/s, and injection was performed at a rate of 100 nl/min. In total, 200-300 nl was injected in one site. The capillary was left in place for 5 min following the injection and then withdrawn at a rate of 0.1 mm/s. For analgesia, buprenorphine (0.05-0.1 mg/kg) was given subcutaneously postoperatively and twice daily for the next 2 days. Experiments started 3-6 weeks after surgery.

For manipulations with the activity of CaMKIIa-RNs, we used the chemogenetic approach. For activation and inactivation of CaMKIIa-RNs in the caudal medulla (the area of the medullary reticular nucleus ventral part and the intermediate reticular nucleus, MdV-IRt area), respectively, AAV5-CaMKIIa-hM3Dq-mCherry-WPRE and AAV5-CaMKIIa-hM4Di-mCherry-WPRE (5 × 10^12^ vg/ ml, Viral Vector Facility, University of Zurich, v.96 and v.102) were injected in wild type mice unilaterally with the following coordinates: - 7.4 mm antero-posterior from bregma, 0.7 mm lateral, and 5.6 mm ventral. To confirm specificity of effects caused by manipulation with activity of CaMKIIa-RNs in this area, AAV5-hsyn-lox-hM3Dq-mCherry-lox-WPRE (4 × 10^12^ vg/ml, Viral Vector Facility, University of Zurich, v.89) was unilaterally injected to the same area in Vglut2-Cre and GAD67-Cre mice to target all glutamatergic and GABAergic neurons, respectively. To check that the effects of manipulation with activity of CaMKIIa-RNs were specific for MdV-IRt area, AAV5-CaMKIIa-hM3Dq-mCherry-WPRE was unilaterally injected in rostral and dorsal adjacent areas in wildtype mice (Fig. 6). The following coordinates for injections to the adjacent areas were used: -6 mm antero-posterior from bregma, 0.7 mm lateral and 5.6 mm ventral, for targeting the dorsal adjacent area; -7.4 mm antero-posterior from bregma, 0.7 mm lateral, and 4.5 mm ventral for targeting the rostral adjacent area. To identify the neurotransmitter type of CaMKIIa-RNs, labelling of CaMKIIa-RNs was combined with RNAscope. For this purpose, we injected 300 nl AAV5-CaMKIIa-EGFP-Cre-WPRE (6 × 10^12^ vg/ml, Viral Vector Facility, University of Zurich, v.315) into wild type mice.

### Experimental designs

For activation/inactivation of infected neurons, CNO (Tocris, catalog no. 4936) dissolved in DMSO was administered intraperitoneally at a dose of 1 mg/kg. Mice performed each of four basic motor behaviors (standing on a horizontal surface, postural corrections on a tilting platform, forward locomotion, and righting) before and between 40-90 min after injection of CNO. Experimental designs used for the behavioral experiments were described earlier (Vemula et al., 2019; Zelenin et al., 2021) and are presented here in brief.

To analyze the basic body orientation and configuration during *standing on a horizontal surface,* the animal was positioned on a transparent tilting platform (12 × 12 cm) that was oriented horizontally (Fig. 2**A**,**D**). To evoke *postural corrections*, the platform with the animal, whose sagittal plane was aligned to the axis of the platform rotation, was tilted periodically in the frontal (transverse) plane of the animal (roll tilt α) with the amplitude of ±20° (Fig. 3**A**,**B**). A trapezoid tilt trajectory with the transitions between extreme positions lasting for ∼0.5–1 s, and each position maintained for ∼1–1.5 s was used. It was necessary to habituate animals to the tilting platform and to train them to stand still during tilts. For this purpose, the animal was positioned on the tilting platform, and tilts with increasing amplitude were applied. If the animal started to walk, it was returned to the initial standing position by the experimenter. Usually, a 20 min session of such training performed during 2–3 days was sufficient to evoke episodes, in which the mouse maintained the standing posture with its sagittal plane aligned to the axis of the platform rotation leading to generation of postural corrections in response to 5-7 sequential tilt cycles.

*Forward locomotion* was performed in a tunnel setup. The setup consisted of a tunnel (length 50 cm, height 4 cm, width 2.5-3.5 cm) with a small box (7 × 7 × 4 cm) at each end of the tunnel. Each box had a removable top and a door that closed the entrance to the tunnel. The animal was placed in the entrance box through the removable top, the top was closed, then the doors to the tunnel of both boxes were opened and the animal could easily walk in the tunnel straight forward but could not turn around. When the mouse entered the box on another side of the tunnel, it turned and performed forward locomotion in the opposite direction. Usually, the animal spontaneously exhibited 3-4 sequential episodes of forward locomotion. Each episode of forward walking in the tunnel consisted of 5-9 steps.

To evoke *righting behavior* (Fig. 5**A**), a mouse was positioned on its back (with its ventral side up) on a horizontal surface. Then the animal was released so it could assume the normal body orientation characteristic for standing.

### Recording and data analysis

The hindlimbs and trunk of the animal were shaved, and markers were drawn on the skinalong the spine. The video camera was positioned at a distance of ∼2 m from the mouse. To characterize the kinematics of fast movements performed during locomotion and righting, high-speed video recording (100 frames/s) was used. Standing on a horizontal surface as well as relatively slow postural corrections were video recorded with a lower speed (50 frames/s). We recorded the view from above during locomotion, the side view during righting, the rear/front view during standing on a horizontal platform, and the rear view during postural corrections on the tilting platform. During standing, simultaneously with recording of the rear/front view, the view from below was recorded (by means of a 45° tilted mirror positioned under the transparent surface). The video recordings were analyzed off-line frame by frame.

In the following text, terms “ipsilateral” and “contralateral” indicate the side of the mouse, respectively, ipsilateral and contralateral to the side of the virus injection.

The head and trunk orientation in the transverse plane (the roll tilts) were characterized by the angles between the vertical (solid yellow lines in Figs. 2**A,D**, 3**A,B**) and the dorso-ventral axis of the head or the trunk, correspondingly (red dashed lines in Figs. 2**A,D**, 3**A,B**). The dorso-ventral axis of the trunk was estimated from the rear view as a line connecting the marker on the spine rostral to the pelvis with the base of the tail. The dorso-ventral axis of the head was estimated from the frontal view as a line perpendicular to the line connecting the eyes. The ipsilateral and contralateral roll tilt angles had positive and negative values, respectively.

Asymmetry in extension/flexion of the hindlimbs during standing on the horizontal surface and during postural corrections was estimated from the rear view and characterized by the extension/flexion asymmetry index: *I_E/F_* = (*L_CONTRA_ - L_IPSI_*)/(*L_CONTRA_* + *L_IPSI_*), where *L_IPSI_* and *L_CONTRA_* were, respectively, the ipsilateral and the contralateral limb lengths. The limb length was estimated by the distance from the heel to the marker on the spine rostral to pelvis (illustrated in Fig. 2**A**, 3**F**). Thus, *I_E/F_* = 0 if the lengths of the left and right limbs were equal, *I_E/F_* > 0 if the contralateral limb was more extended than the ipsilateral one, and *I_E/F_* < 0 if the ipsilateral limb was more extended than the contralateral one.

To characterize the position of the limbs in relation to the trunk (abduction/adduction) during standing on horizontal surface, the view from below was used. The trunk outline and its midline were drawn. Then the axes perpendicular to the midline at the level of forelimbs (one-third of the distance from the nose to the base of the tail) and hindlimbs (two-thirds of the distance from the nose to the base of the tail) were drawn. The midpoint of the body width at the corresponding level was taken as “0”. The position of a limb in relation to the trunk was characterized by the coordinate along the corresponding axis (*b* in Fig. 2**G**). This coordinate was termed “the lateral position” of the limb and expressed in percent of the body half-width. Positive and negative values of the lateral position of the limb indicated ipsilateral and contralateral location of the foot in relation to the midline, respectively (Fig. 2**G-I**).

Asymmetry in abduction/adduction of the fore- and hindlimbs was characterized by the abduction/adduction asymmetry index: *I_AB/AD_* = (*b_CONTRA_ - b_IPSI_*)/(*b_CONTRA_ + b_IPSI_*), where *b_CONTRA_* and *b_IPSI_* were the lateral positions of the contralateral and the ipsilateral limbs, respectively. Thus, *I_AB/AD_* = 0 if the lateral positions of the left and right limbs were symmetrical in relation to the trunk, *I_AB/AD_* > 0 if the contralateral limb was more abducted than the ipsilateral one, and *I_AB/AD_* < 0 if the ipsilateral limb was more abducted than the contralateral one.

To estimate the trunk orientation in the transverse plane stabilized on the tilting platform, the rear view was used. We measured the roll tilt angle of the trunk at two conditions: when the animal was standing on the platform tilted to the right and to the left (shown by black and gray in Fig. 3**C**). The average of these two angles was termed “the stabilized angle” [the angle between the vertical (green line) and the dashed red line in Fig. 3**C**]. With perfect stabilization of the dorsal side-up trunk orientation, the stabilized angle is equal to zero (Fig. 3**C**, left panel), while positive and negative values of the stabilized angle indicate stabilization of orientation with the ipsilateral and contralateral roll tilt, respectively. To characterize the efficacy of postural corrections, we calculated the coefficient of postural stabilization *K_STAB_* = 1 – β/α, where β is the amplitude of the trunk roll tilt, and α is the amplitude of the platform tilt (Fig. 3**I**). With perfect stabilization, *K_STAB_* = 1; with no stabilization, *K_STAB_* = 0.

To characterize the trunk orientation in the transverse plane maintained during locomotion, the position of the spine in relation to the left-right body edges was characterized by using the top view. The trunk outline and its midline were drawn. Then the axis perpendicular to the midline at the level of hindlimbs (two-thirds of the distance from the nose to the base of the tail) was drawn. The midpoint of the body width was taken as “0”. We measured the deviation of the spine (the red point in Fig. **4A-C**) from the midpoint of the body width (“0”) to the right (at the moment of the left hindlimb lift-off; *a* in Fig. 4**A**) and to the left (at the moment of the right hindlimb lift-off) and calculated the average of these two values. This average was termed “the spine position during locomotion” and expressed in percent of the body half-width.

Asymmetry in abduction/adduction of the hindlimbs during locomotion was characterized by the abduction/adduction asymmetry index: *I_AB/AD_* = (*b_CONTRA_* – *b_IPSI_*)/(*b_CONTRA_* + *b_IPSI_*), where *b_CONTRA_* and *b_IPSI_* were the lateral positions of the ipsilateral and the contralateral limbs at the corresponding lift-off moments (when the maximal lateral displacement of the spine was observed, Fig. 4**A**).

### Tissue immunochemistry

Mice were euthanized by anesthetic overdose with pentobarbital (250 mg/kg), and perfused transcardially with 4°C saline followed by 4% paraformaldehyde. Brain and spinal cord tissue was dissected free and then postfixed in 4% paraformaldehyde for 3 h at 4°C. Tissue was cryoprotected by incubation in 15% sucrose in phosphate-buffered saline (PBS) overnight. Tissue was then embedded in Neg-50 medium (Thermo Fisher Scientific) for cryostat sectioning. Coronal sections were obtained on a cryostat and mounted on Superfrost Plus slides (Thermo Fisher Scientific). Brainstem and spinal cord coronal sections were cut at 40 µm thickness.

For immunohistochemical detection of mCherry, sections were incubated overnight with a rabbit anti-DsRed/tdTomato/mCherry (1:1,000, Clontech, catalog no. 632496). Both the primary and the secondary antibodies were diluted in 1% bovine serum albumin (BSA), 0.3% Triton X-100 in 0.1 M PB. Following incubation with the primary antibody over night at 4°C, the sections were incubated for 3 hr with secondary antibody: Alexa-568 anti-rabbit (1:500; Jackson Immuoresearch). Then, slides were washed in PBS-T, counterstained with Hoechst 33342 (1:2,000) and mounted with coverslips using glycerol containing 2.5% diazabicyclooctane (DABCO; Sigma-Aldrich). Sections were imaged using either a Zeiss widefield epifluorescence microscope or a Zeiss LSM 800 confocal microscope.

### Heatmap generation

To build heatmaps showing the extent of the infected area in the brainstem and areas of the spinal cord gray matter with terminals of the infected neurons, we used images of coronal sections of the brainstem and spinal cord (cervical segments C4-C6, thoracic segments T5-T9, and lumbar segments L2-L5) containing the infected areas. Four borders of an image were aligned with the section edges. The image area was divided into 500 grid regions of interest (ROIs) by 25 columns and 20 rows. The fluorescence intensity *F* was measured in each ROI, the maximal *F* across all 500 ROIs was found, and *F* was normalized to this maximum. To obtain an averaged heatmap for a particular brainstem or spinal cord level, we used heatmaps built for individual sections taken at this particular level from different animals, and for each ROI we averaged the normalized F values across this set of heatmaps.

### RNAscope *in situ* hybridization

For *in situ* hybridization (RNAscope), the tissue sections were processed using RNAscope Multiplex Fluorescent v2 Assay (Advanced Cell Diagnostics, ACD, 323110) according to the manufacturer’s instructions. In brief, the sections were air dried for 1 h at room temperature (RT) and placed in the PBS for 5 mins then dehydrated through sequential EtOH steps (50%, 70%, 100%, 100%) before air drying at RT. Then the sections were treated with hydrogen peroxide for 10 min at RT, washed in water, and treated with target retrieval reagents for 5 min at 99°C. Sections were rinsed with water and treated with 100% EtOH for 3 min at RT and air dry again at RT. Tissues were then treated with protease III for 30 min at 40°C before being rinsed with water. Specific probes were hybridized with sections for 2 h at 40°C in a humidified oven, rinsed in wash buffer, and then stored overnight in 5× saline sodium citrate (SSC) buffer. The next day, the sections underwent a series of AMP1∼3 incubation steps to amplify and develop the signals. The corresponding HRP channels and fluorophores (TSA Plus Fluorophores, Akoya) were applied to develop the signals from the hybridized probes. The following probes were used: Mm-Slc17a2-C1 (#319191-C1), Mm-Slc32a1-C1 (#456751-C1), and Mm-CamkIIa-C2 (#445231-C2). After finishing all the RNAscope steps, the sections were applied to the immunohistochemistry with the RFP antibody and Nissl.

### Statistical analysis

Sample size was not estimated *a priori* to obtain a given power. Mice were randomly allocated to different groups for the in *vivo* experiments using a block design. Data sampling and analysis were not blinded. All quantitative data in this study are presented as the mean ± SD. Mean values were calculated as averages of the mean from each animal. Each parameter was measured in an individual animal 8-80 times. The Student’s *t* test (two-tailed) was used to characterize statistical significance when comparing different means. Spearman’s rank correlation coefficient was calculated to correlate the changes in the duration of the righting reflex movement with the asymmetry indexes. The significance level was set at *p* = 0.05.

## AUTHOR CONTRIBUTIONS STATEMENT

L.-J.H., P.Z., and. T.D. designed the study. L.-J.H., and V.L. performed histological analysis and tracings. L.-J.H., P.Z., and V.L. performed behavioral experiments and analyzed data. S.- H.C. and F.L. performed RNAscope and provided related resources. T.D. provided the funding and technical support for the work. L.-J.H. and T.D. wrote the paper with contributions from all authors. All authors approved the final version of the manuscript.

## CRediT authorship contribution statement

**Pavel V. Zelenin:** Conceptualization, Investigation, Methodology, Writing – review and editing. **Vladmir F. Lyalka:** Investigation, Writing – review and editing. **Shih-Hsin Chang:** Methodology, Writing – review and editing. **Francois Lallemend:** Methodology, Funding acquisition, Writing – review and editing. **Tatiana G. Deliagina:** Conceptualization, Funding acquisition, Investigation, Methodology, Validation, Visualization, Writing – original draft. **Li-Ju Hsu:** Conceptualization, Formal analysis, Investigation, Methodology, Validation, Visualization, Writing – original draft.

## Declaration of competing interest

The authors declare that they have no competing interests.

## Data availability

The data that support the findings of this study are available from the corresponding author upon request.

## Acknowledgements

This work was supported by grant from NIH (R01 NS-064964) to TGD; by grants from Swedish Research Council (2020-02502) to TGD; KI fonder (2024-2026) to LJH; Alice and Wallenberg foundation (KAW2022-0130) to FL; Olle Engkvist foundation (228-0277) to FL; NSTC postdoctoral research abroad program from Taiwan (112-2917-I-564-029) to SHC.

## Notes

### Competing Interest Statement

The authors have declared no competing interest.

